# coil: an R package for cytochrome C oxidase I (COI) DNA barcode data cleaning, translation, and error evaluation

**DOI:** 10.1101/2019.12.12.865014

**Authors:** Cameron M. Nugent, Tyler A. Elliott, Sujeevan Ratnasingham, Sarah J. Adamowicz

**Affiliations:** Department of Integrative Biology, University of Guelph. Guelph, Ontario, Canada; Centre for Biodiversity Genomics, Biodiversity Institute of Ontario, University of Guelph. Guelph, Ontario, Canada

**Keywords:** COI, DNA barcoding, error identification, translation

## Abstract

Biological conclusions based on DNA barcoding and metabarcoding analyses can be strongly influenced by the methods utilized for data generation and curation, leading to varying levels of success in the separation of biological variation from experimental error. The five-prime region of cytochrome c oxidase subunit I (COI-5P) is the most common barcode gene for animals, with conserved structure and function that allows for biologically informed error identification. Here, we present coil (https://CRAN.R-project.org/package=coil), an R package for the pre-processing and error assessment of COI-5P animal barcode and metabarcode sequence data. The package contains functions for placement of barcodes into a common reading frame, accurate translation of sequences to amino acids, and highlighting insertion and deletion errors. The analysis of 10,000 barcode sequences of varying quality demonstrated how coil can place barcode sequences in reading frame and distinguish sequences containing indel errors from error-free sequences with greater than 97.5% accuracy. Package limitations were tested through the analysis of COI-5P sequences from the plant and fungal kingdoms as well as the analysis of potential contaminants: nuclear mitochondrial pseudogenes and *Wolbachia* COI-5P sequences. Results demonstrated that coil is a strong technical error identification method but is not reliable for detecting all biological contaminants.

## Introduction

DNA barcoding leverages sequence diversity within standardized gene regions for the identification and classification of organisms (Hebert *et al.* 2003; Ratnasingham and Hebert 2007). Answering questions about biodiversity through DNA barcode analyses depends on the comparison of novel barcode sequences to reference libraries or the *de novo* comparison of sequences to one another (Hebert *et al.* 2004; Ratnasingham and Hebert 2007; Hubert and Hanner 2015; Elbrecht *et al.* 2018). Techniques such as DNA metabarcoding greatly expand the complexity of comparative analyses due to the increased scale and associated challenges such as additional noise in datasets (Cristescu 2014). Barcode and metabarcode output sequences can vary in terms of both length and accuracy due to a mixture of true biological variation (Pentinsaari *et al.* 2016), the primers or sequencing platforms utilized (Folmer *et al.* 1994; Hebert *et al.* 2018), and the data cleaning steps employed (Elbrecht *et al.* 2018). Defining the boundaries of the barcode region, trimming adjacent sequence information, and aligning barcode information in a common context makes barcode sequences more directly comparable, thereby allowing for more accurate analysis of the evolutionary history and relatedness of organisms.

An approximately 657bp fragment of the 5’ region of the cytochrome C oxidase subunit I gene (COI-5P) is the main marker utilized in DNA barcoding of the animal kingdom (Hebert *et al.* 2003). COI is a component of the last enzyme in the electron transport chain, which is essential to metabolism (Castresana *et al.* 1994; Pentinsaari *et al.* 2016). As a result, the COI-5P barcode region has particular selective constraints, especially against changes to the corresponding amino acid sequence that modify the protein structure and negatively affect metabolism (Tsukihara *et al.* 1995; Pentinsaari *et al.* 2016). The conserved amino acid sequence corresponding to the COI-5P barcode region means that translation of sequences can provide a powerful means of detecting insertion and deletion technical errors (indels) that have resulted in reading frame shifts that alter the inferred amino acid profile.

The wealth of COI-5P sequence data available and the conserved profile of the sequence provide the information necessary for pre-processing and evaluation of novel COI-5P barcode data (Ratnasingham and Hebert 2007; Stoeckle and Kerr 2012; Pentinsaari *et al.* 2016; Porter and Hajibabaei 2018). At present, the comparison and evaluation of sequences in a common context can be accomplished through the *de novo* alignment of novel sequences, but this approach grows in computational complexity with the volume of data and the rate of errors in the data (Elbrecht *et al.* 2018). Software tools that allow for the local or web-based alignment of novel sequences to reference datasets include MACSE (Multiple Alignment of Coding SEquences) (Ranwez *et al.* 2011) and USEARCH (Edgar 2010). Analysis of novel sequences with these methods relies on the comparison of inputs against large sets of reference sequences which a user must acquire, store, and curate. The outputs must also be scrutinized in subsequent analysis so that the presence of errors can be inferred from deviations in alignment. Additionally, if reference data sets are not sufficiently cleaned, then sequences with errors may not be identified, and if references contain taxonomic bias then true biological variants may be interpreted as errors. If systematic errors exist in novel samples, for reasons such as the tendency of a given sequencing platform more often to insert or delete bases in certain regions of a sequence (Hebert *et al.* 2018) or due to bias related to certain sequence motifs (Schirmer *et al.* 2016), then *de novo* alignment of data may not identify systematic errors and instead depict these as biological variants. Another method for processing and analyzing barcode sequences is to upload them to a public repository such as BOLD (boldsystems.org), which uses proprietary error evaluation software (Ratnasingham and Hebert 2007). If the analyses being conducted are exploratory or still in the development stage, then a researcher may not yet be prepared to upload the data to a public repository. Additionally, if methods and associated results are being refined, the repeated upload of sequences to BOLD for processing and error checking can be inefficient and best left as a final step in the workflow. A locally available tool for the establishment of reading frame for COI-5P data could allow researchers to compare barcode data from different samples efficiently in a common context. Additionally, placing DNA sequences in the correct reading frame permits the translation of the nucleotide sequence to amino acids, thereby providing a means of indel identification. Insertions and deletions that introduce frameshifts can be easily detected through the resulting drastic changes to the amino acid sequence.

A tool for pre-processing and technical error identification in DNA barcodes would be a useful addition to both DNA barcode and metabarcode workflows. In DNA barcoding, the barcode sequences generated for a set of specimens can be evaluated and data correction efforts quantitatively targeted. Barcodes displaying evidence of indel errors could be excluded, subjected to error correction, or recommended for resequencing. In DNA metabarcoding, the haplotypes contained within identified operational taxonomic units (OTUs) could be processed and tested for evidence of indels. Generation of consensus sequences for OTUs using only sequences without evidence of technical errors would improve quality. Additionally, if intraspecific diversity were being characterized through the examination of haplotypes within OTUs, low-abundance haplotypes would not have to be excluded solely based on their abundance (Elbrecht *et al.* 2018) and could be included in subsequent analyses with higher confidence if they displayed no evidence of technical errors.

Here, we present a new R package for the pre-processing of COI-5P DNA barcode data: coil. The package provides several functions for: (1) placement of sequences in a common reading frame, trimming of sequences to the COI-5P barcode region, (3) translation of nucleotides to amino acid sequences, and (4) estimating the likelihood that a sequence contains an insertion or deletion error. The tool is free and publicly available through CRAN (https://CRAN.R-project.org/package=coil) and GitHub (https://github.com/CNuge/coil). We demonstrate the effectiveness of coil by showing how it can be used to align novel barcode sequences to the COI-5P profile and to identify sequences with insertion or deletion errors, in most cases with greater than 97.5% accuracy.

## Implementation

The coil package contains functions that allow new DNA barcode data to be placed into reading frame, translated, and assessed for evidence of insertion or deletion errors (Fig. 1). The statistical models the package uses to evaluate sequences are two profile hidden Markov models (PHMMs) respectively trained on nucleotide and amino acid sequences of the COI-5P barcode region. Profile hidden Markov models (PHMMs) are a specialized variant of a hidden Markov model well suited for statistically representing biological sequences that share a common evolutionary origin (Durbin and Eddy 1998; Eddy 1998; Wilkinson 2019). A PHMM is a probabilistic representation (profile) of a multiple sequence alignment. Training a PHMM captures information about the structure of a series of related biological sequences, characterizing the probability of an observed character being present at the given point in the sequence and the probability of transitioning from the observed character to the next character in the sequence. Once trained, PHMMs are probabilistic representations of the aligned training sequences that can be used to determine how closely new sequences match the profile. The characters in the new biological sequence (the nucleotide or amino acid alphabets) constitute the observed characters (states), and comparison to the PHMM relates the observed characters to the series of unknown hidden states that describes how the observed sequence of characters most likely match to the probabilistic profile (Figs. 2, 3). The PHMMs used within coil are meant to take a new DNA barcode sequence (the observed states) and look for evidence of erroneously inserted or deleted base pairs in the sequence (the hidden states). For coil, the observed character states are both the nucleotides and amino acids of the barcode sequence and the hidden states are whether the given character: (1) matches the barcode profile, (2) is likely an erroneous inserted character, or (3) is evidence of a missing (deleted) character.

**Fig. 1.**
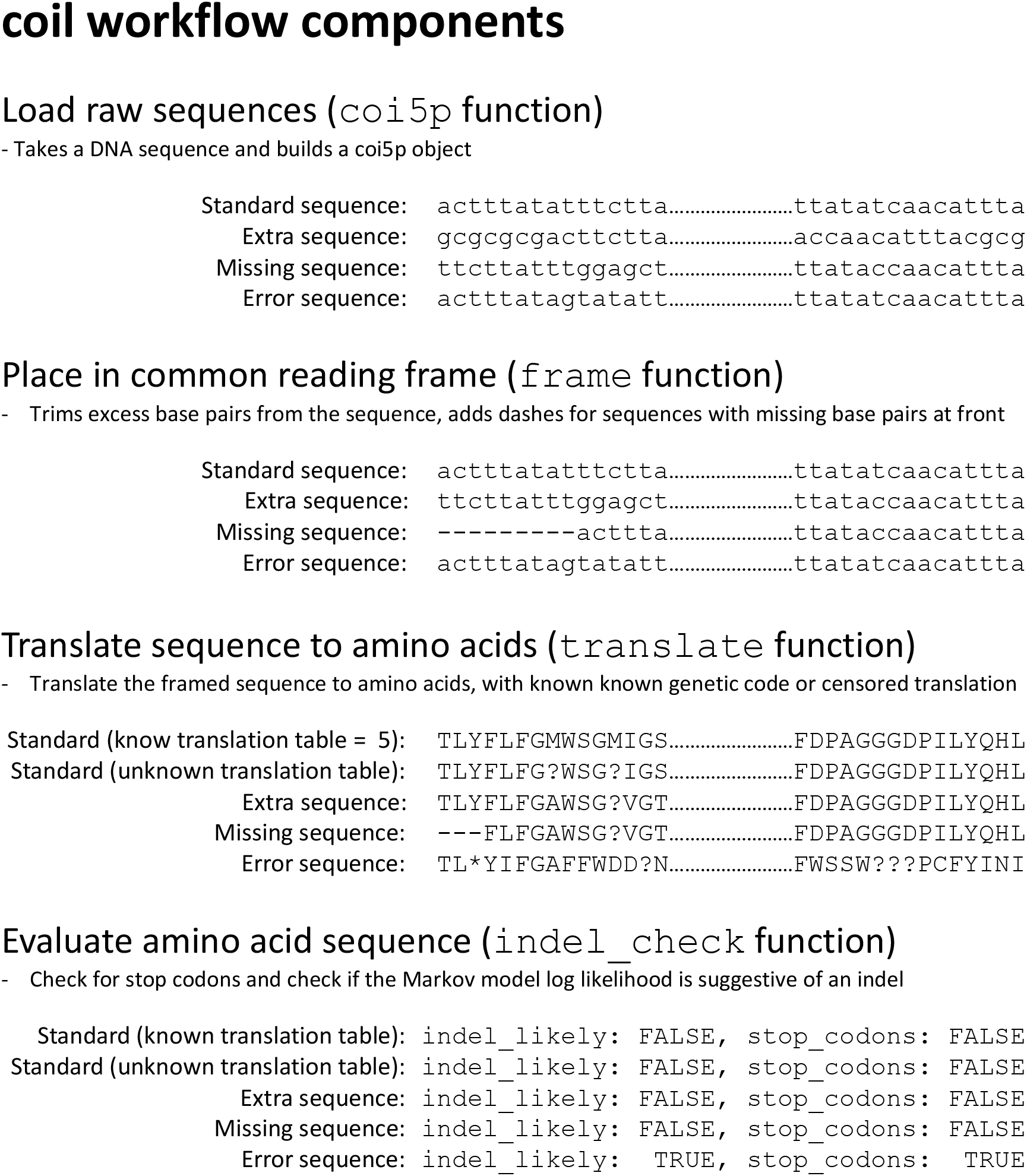
Depiction of the coil library’s analysis pipeline. The tasks performed by the four main functions of the analysis pipeline on a variety of potential use cases are shown, and a brief explanation is given. For information on how to implement the analysis pipeline, please consult the package’s vignette (Supplementary File 7).

**Fig. 2.**
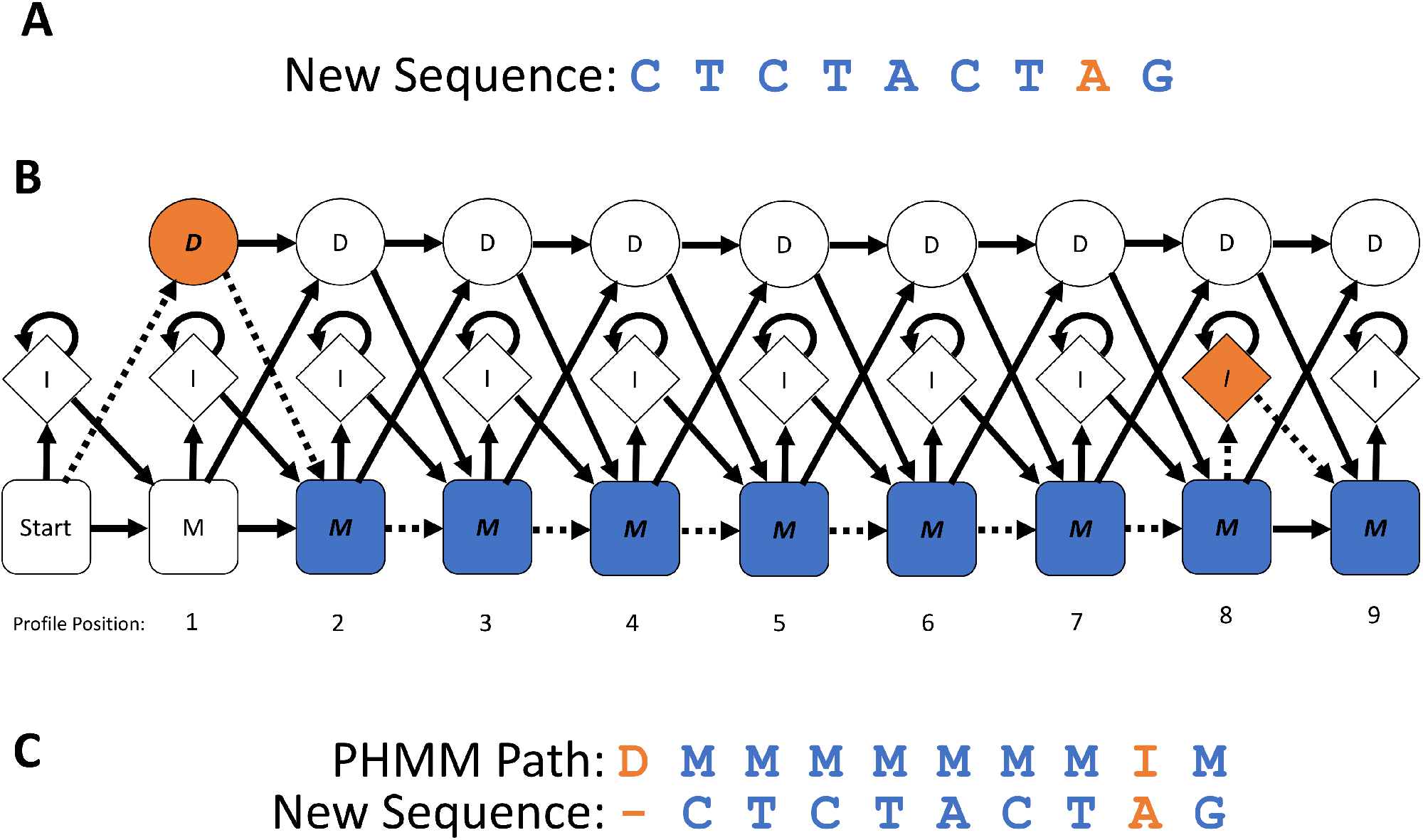
Description of how a trained profile hidden Markov model (PHMM) determines the hidden state of nucleotides for a new biological sequence. The input (A) is a truncated example of a DNA sequence (the nucleotides are the observed states) to be evaluated using the PHMM. An erroneous nucleotide (indicated in orange) has been inserted into the sequence; this fact would be unknown prior to evaluation. (B) A diagram of the potential paths of hidden states through the trained PHMM (only first 9 of 657 profile positions shown). The possible hidden states are: M (indicates a nucleotide is a match to the sequence profile), D (indicates the nucleotide for the given position in the profile has been deleted), and I (indicates a nucleotide has been inserted into the sequence at that position). The arrows indicate the possible transitions from one hidden state to another, and the numbers below the diagram indicate the positions in the COI-5P profile. The Viterbi algorithm is used to compare the observed states (nucleotide sequence) to the PHMM to determine the most probable set of hidden states corresponding to the sequence. When a DNA sequence is evaluated against the PHMM using the Viterbi algorithm, the position-specific emission and transition probabilities (essentially the probability of a given nucleotide being found at that point in the sequence and the probability of transitioning from the current nucleotide to the next observed nucleotide, see Fig. 3) are used to assign hidden states to the nucleotides in the sequence. The most probable hidden state path aligning the new sequence to the displayed section of the PHMM is indicated by the series of coloured hidden state cells (with italicized letters) and the dotted arrows indicating the transitions between states. (C) In this example, a nucleotide was missing at the start of the sequence, so the hidden state for position 1 in the profile is delete (‘D’). The orange (inserted) nucleotide has been identified as an insertion between positions 8 and 9 of the profile, and therefore this nucleotide is assigned the hidden state of insert (‘I’), while all other nucleotides are found to match positions in the profile and have been assigned hidden states of match (‘M’).

**Fig. 3.**
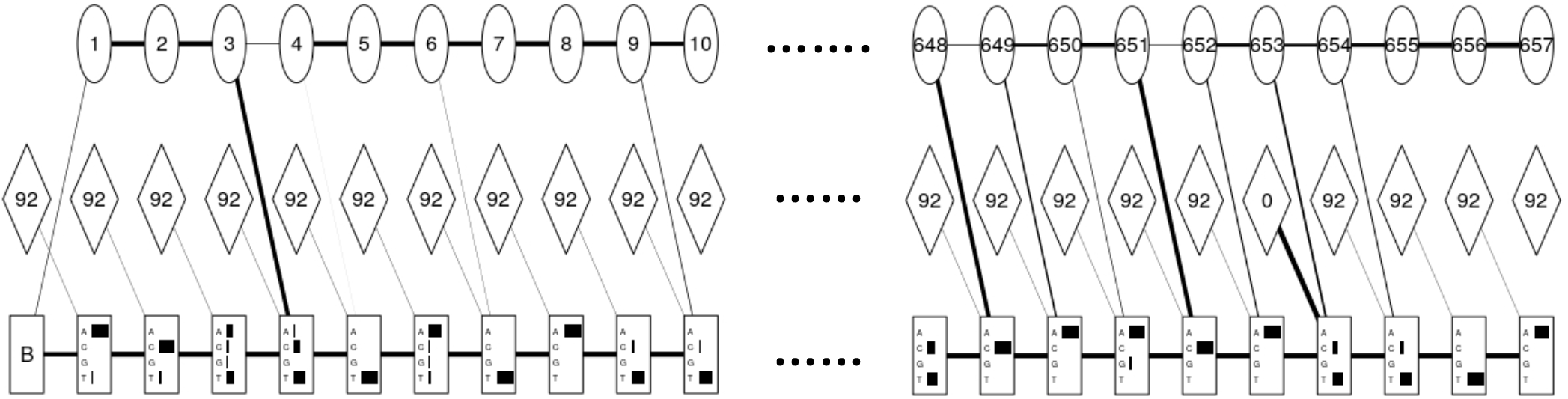
Diagram of the emission and transition probabilities for the first and last 10 positions in the nucleotide PHMM used in the coil package. The circles represent the hidden delete states and contain the profile positions, the diamonds indicate the hidden insert states, and the numbers within the diamonds indicate the probability of insert-insert transitions. The rectangles indicate the match states, and the horizontal bars within represent the emission probabilities (synonymous with nucleotide frequencies) for the possible nucleotides at the given profile position. The weight of the lines between shapes represent the transition probabilities.

The coil analysis pipeline establishes the reading frame of sequences (Fig. 4) using the Viterbi algorithm (Fig. 1) to compare the series of observed states (the DNA sequence) to the nucleotide PHMM (Durbin and Eddy 1998; Wilkinson 2019). This algorithm determines the most likely sequence of hidden states corresponding to the observed sequence by determining the optimal path through the network of emission and transition probabilities, which are respectively defined as the probability of a given base at a given position and the probability of a given base following the preceding base at the given position. For framing of nucleotide data, the output is used to infer the presence of DNA not matching the 657bp region defined by the boundaries of the model or missing information at the start of the sequence (Figs. 1, 4). It is important to note that any data outside of the 657bp Folmer region (upon which the model is trained) is removed and not utilized in subsequent translation or error assessment; coil reports when information is trimmed from the input sequence as part of this processing step.

**Fig. 4.**
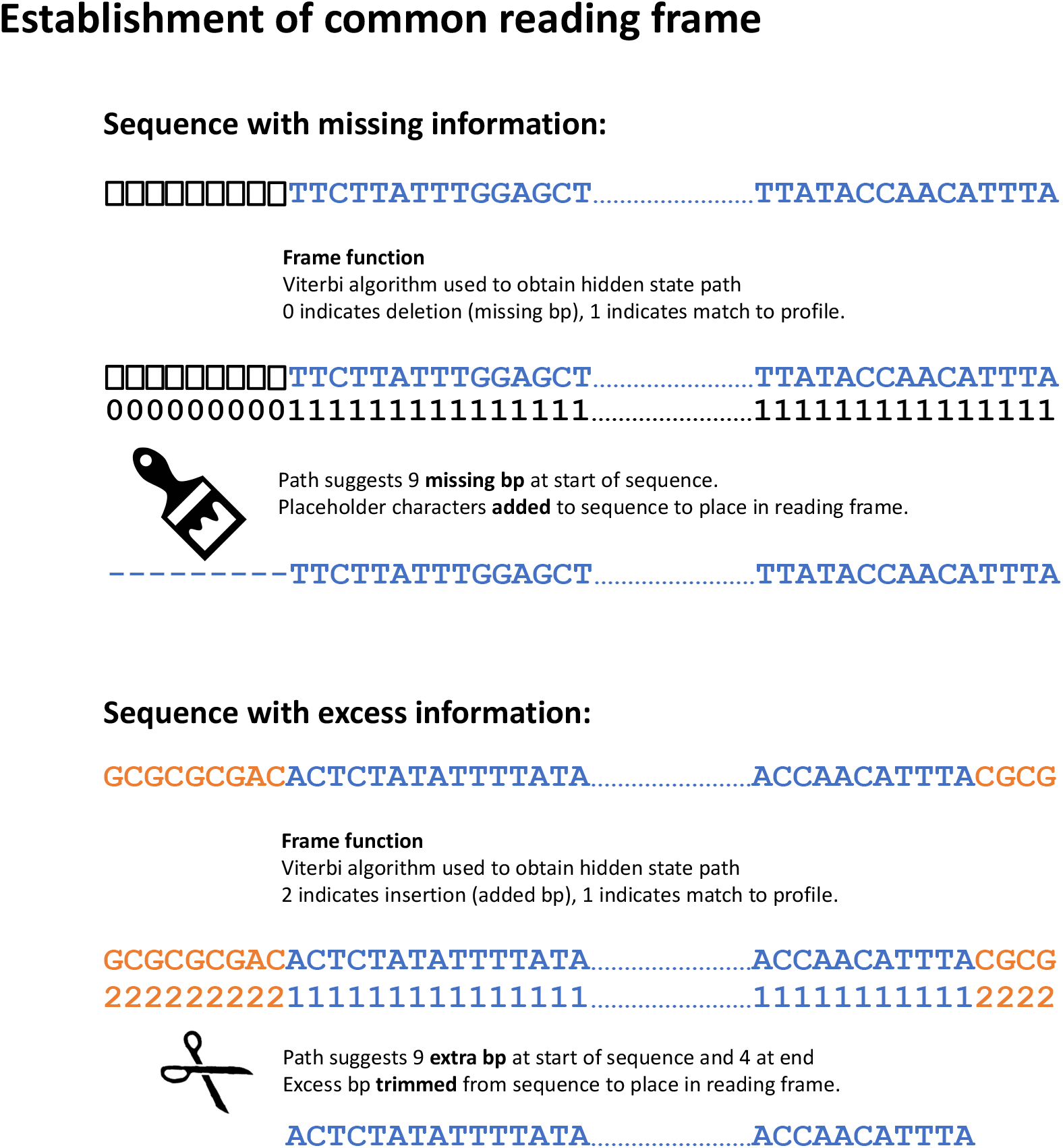
Diagram showing how coil establishes the reading frame of nucleotide sequences through assessment of the hidden state path produced through the Viterbi algorithm’s comparison of input sequence to the nucleotide PHMM.

### Translation

As a result of its ancient endosymbiotic origin, mitochondrial translation utilizes a different genetic code from the nucleus, and this code additionally varies across taxa (Youle 2019). The coil package is therefore designed in a manner robust to the variability in animal mitochondrial genetic codes (NCBI numeric identifiers: 2, 4, 5, 9, 13, 14, 21, 24, and 33) (Elzanowski and Ostell 2019). If the taxonomic designation of a barcode sample is known and provided along with the input sequence, then coil calls functions from the package SeqinR (https://CRAN.R-project.org/package=seqinr) to translate the sequence using the appropriate genetic code (Osawa *et al.* 1992; Jukes and Opsawa 1993; Charif and Lobry 2007). By default, a special censored translation option is utilized by coil in order to accommodate instances when the taxonomy of a sample (and therefore the correct genetic code) is unknown. Codons that vary in the amino acid they code for across known animal mitochondrial genetic codes are translated to a placeholder character (‘?’) to indicate that the amino acid at this location in the sequence cannot be stated with certainty (Fig. 5). This allows coil to assess the likelihood of the sequence containing indel errors, without being biased by errors introduced due to the appropriation of the wrong genetic code. As an example of why this is necessary, if a barcode sequence from a biting fly (genetic code = 5) was amplified in the analysis of a mammal host specimen (proper genetic code = 2) (Ratnasingham and Hebert 2007; Elzanowski and Ostell 2019) then the codon ‘AGG’ would be interpreted as a stop codon, as opposed to the correct amino acid: serine. The presence of a stop codon due to this translation error would make the resulting amino acid sequence highly improbable and lead to it being flagged as likely erroneous, when it is in fact a true barcode sequence that a user may in some instances wish to retain. To avoid this, censored translation outputs placeholder characters that do not negatively impact the likelihood of the amino acid sequence because of mistranslated codons. The censored translation outputs do, however, lead to a slight loss of power for subsequent error assessment due to reduced sample size because placeholder characters (for 5/64 codons) are not evaluated in the calculation of the likelihood score.

**Fig. 5.**
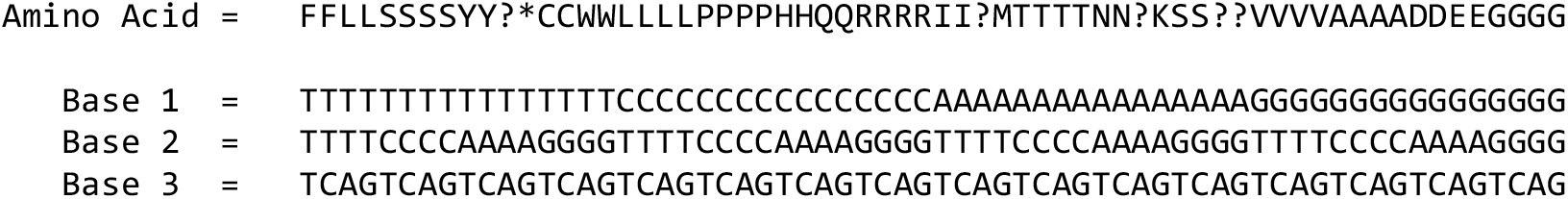
The censored genetic code utilized by coil to translate codons of nucleotides to amino acids when the genetic code corresponding to an input sequence is not known. For each column, the three bottom rows represent the nucleotide composition of the codon (base pair 1, 2, and 3 of the codon read from top to bottom). The amino acid characters in the top row represents the result of translating the codon listed below. The ‘?’ placeholder characters in the Amino Acid row correspond to the ambiguous codons, which code for different amino acids in different animal mitochondrial genetic codes. These amino acids are therefore censored to avoid translation errors, while translation of all other codons is conducted normally.

### Error assessment

The inference of sequence errors is accomplished by comparison of a translated barcode sequence against the amino acid PHMM (Fig. 6). The Viterbi algorithm (Fig. 1) is used to compare the amino acid sequence to the PHMM, and the likelihood of the optimal path that aligns the input sequence to the PHMM is considered. The likelihood is expressed in logarithmic format; the higher a log likelihood (all log likelihood values are negative) the more likely it is that the sequence is a match to the profile. Conversely, the lower the log likelihood, the more likely the amino acid sequence is at some point shifted out of reading frame due to an insertion or deletion error in the nucleotide sequence, leading to a highly unlikely amino acid sequence. Low log likelihood values could result from frameshifts due to technical errors (i.e. PCR amplification or DNA sequencing) or due to the detection of non-barcode sequences, such as nuclear pseudogenes. Amino acid insertions or deletions (indel of a full codon of nucleotides) do not result in frameshifts and lead to only minor perturbations in the log likelihood score; the check of the amino sequence is therefore receptive of mutations of this kind, even if they were unseen in the PHMM training data. The check for frameshifts via the PHMM comparison is complemented by a simple query of the amino acid sequence for stop codons, which can provide additional evidence of frameshift errors or identify instances where translation has been conducted with the wrong genetic code. Since the enzyme that COI-5P codes for is highly conserved, there is an extremely low likelihood of a biologically viable mutation that leads to early termination of the amino acid sequence. The log likelihood of the sequence path and the query of the sequence for stop codons can therefore be used in conjunction to assess sequence validity with high accuracy. This error checking method has the additional advantage of not employing a computationally expensive step of comparison of the input sequence against a large set of references (Edgar 2010; Ranwez *et al.* 2011) and can be utilized without acquisition or user-curation of a large set of reference sequences.

**Fig. 6.**
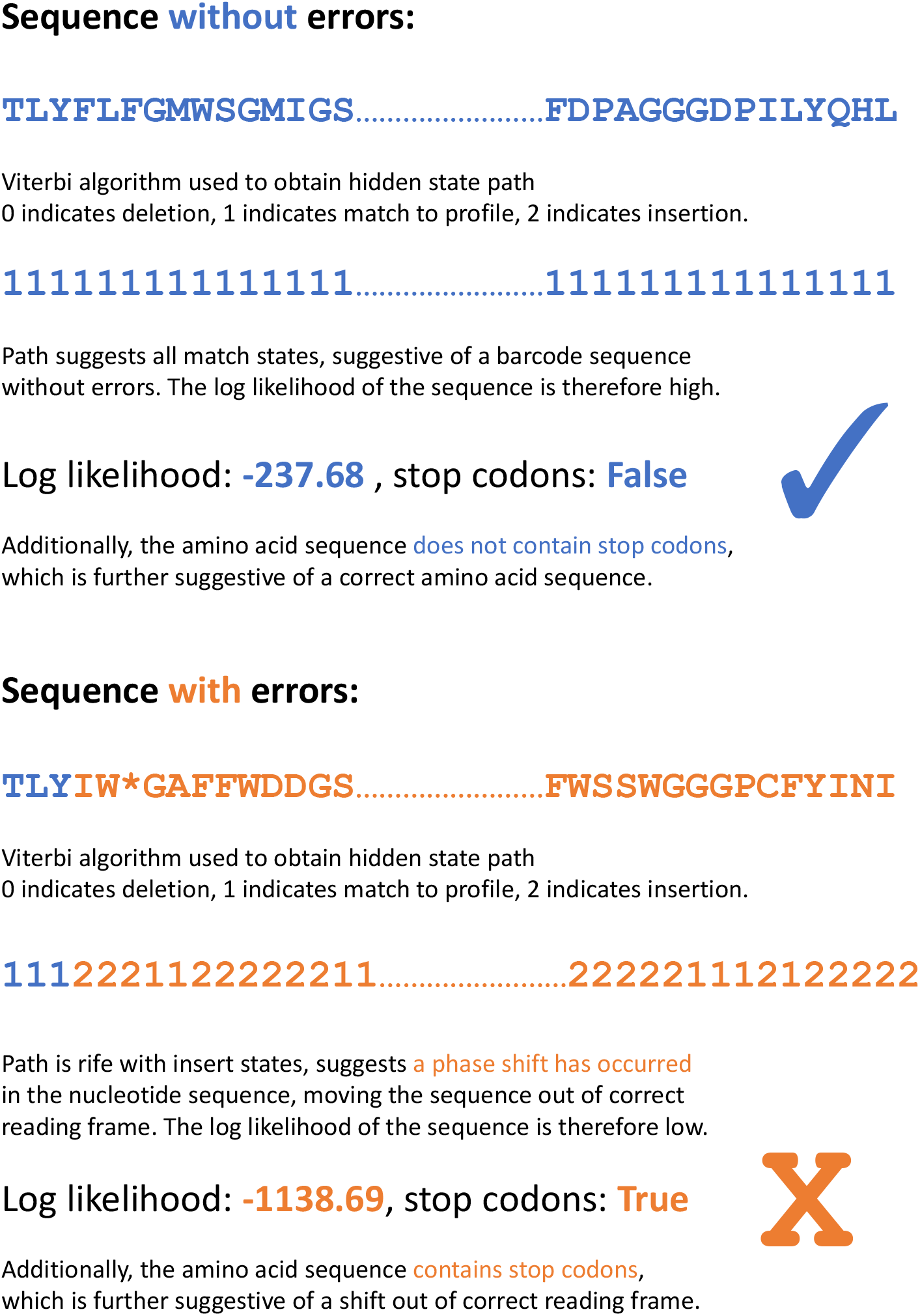
Diagram showing how coil’s indel_check function utilizes the hidden state path output by the Vitertbi algorithm to determine the likelihood that a sequence contains an insertion or deletion error that has led to a shift in reading frame.

## Methods

### BOLD data acquisition

The Barcode of Life Data Systems (BOLD: boldsystems.org) public database was queried to obtain a representative sample of publicly available COI-5P sequences. Unique COI-5P haplotypes (Ratnasingham and Hebert 2013) were retained if the following criteria were met: (1) the sequence was >600bp in length within the Folmer region (this was done to ensure sequences could be aligned with high fidelity prior to model training) (Folmer *et al.* 1994), (2) the taxonomy of the sequence was known to a genus level, (3) there were no missing bp (‘N’) in the Folmer region, (4) the corresponding amino acid sequence did not contain stop codons, (5) the result of BOLD’s internal check for contaminants was negative. The query and subsequent filtering yielded approximately 1.3 million unique sequences (in BOLD’s aligned format). To isolate the 657bp COI-5P region of the sequence, data were initially analyzed in BOLD’s aligned format. Sequences with information outside of the COI-5P region were truncated and the placeholder dashes from BOLD’s aligned format were then removed from the sequences. The resulting unaligned sequence data for only the Folmer region were then used in subsequent model training and assessment. PHMM training (described below) included *de novo* multiple sequence alignment, so that the alignment of input sequences for PHMM generation was repeatable (parameters for BOLD’s alignment of sequences are not publicly available). This process was repeated for the amino acid sequences, with only the 219 amino acids corresponding to the barcode region retained. To ensure that the data used for model training and the data used for model testing and validation were independent from one another, the barcode dataset was first split in a stratified fashion (on the taxonomic level: family) into training (70%), test (15%), and validation (15%) sets for independent use in model training, refinement, and final assessment. The taxonomically stratified split of the data was done to ensure that the training dataset was not biased towards the largest taxonomic orders due to random chance and that subsequent models were trained on diverse data representative of currently-available DNA barcode data on BOLD for the entire animal kingdom. It should be noted that some of the sequences in the BOLD dataset may be taxonomically misidentified, but that any errors of this kind would not have a significant impact on the PHMMs constructed. Training samples were used to train a single nucleotide and a single amino acid PHMM to represent the entire animal kingdom, so misidentified training data would only cause a slight change in the distribution of the taxonomic diversity of the test, validation, and training sets and not likely result in a significant impact on model performance.

### Model training

A nucleotide and amino acid profile hidden Markov model (PHMMs) were respectively trained on COI-5P nucleotide and amino acid training sequences (Supplementary File 1). The models were not trained on the entire set of 936,971 barcodes in the training set; rather, in an attempt to prevent model over fit (and allow effective application on new animal taxa not yet represented on BOLD), different-sized taxonomically representative subsamples of the full set of training data were used to train competing PHMMs. The full training set was subsampled in a stratified fashion (on the taxonomic level: family) so that different taxonomic groups were proportionally represented in the subsamples, with the exception of families with fewer than 10 samples in the training set, which had all their representatives added to each subset to ensure that these rare families were represented in model training. The two taxonomically representative training sets of ~1.2% (11,391 sequences) and ~3.0% (28,189) of the training data were produced, and each was subjected to the following procedure. Training of the nucleotide and amino acid PHMMs was done using the R package aphid (Wilkinson 2019). Model training was conducted using aphid’s derivePHMM function, which produces a *de novo* multiple sequence alignment and generates an optimal PHMM using the Baum-Welch training algorithm (Wilkinson 2019). Training with the derivePHMM function utilized default parameters, except for the ‘pseudocounts’ and ‘maxsize’ parameters. The ‘background’ option was used for the ‘pseudocounts’ parameter (see Durbin and Eddy 1998 – Chapter 5 for explanation of pseudocounts). Initially, models were trained using two ‘maxsize’ parameter pairs: (a) a nucleotide PHMM maxsize of 657 and a corresponding amino acid maxsize of 219 and (b) a nucleotide PHMM maxsize of 673 and a corresponding amino acid maxsize of 224. The ‘maxsize’ parameter limits the upper bound of the number of modules in the PHMM; i.e. it provides the maximum length of the profile that represents the sequences. Alignment of test data to the models via the Viterbi algorithm revealed that when the larger maxsize parameter was utilized, the PHMM expanded to the maximum size permitted. In subsequent testing this led to variable framing of sequences (dependent on taxonomy) and made the establishment of a common reading frame infeasible. The smaller maxsizes of 657/219 yielded a set of PHMMs that could effectively establish the reading frame of taxonomically diverse sequences in the test data, and this pair of PHMMs was therefore retained as the internal PHMMs for coil, against which new barcode sequences are compared. Of the two training data sizes that were used, the PHMMs resulting from training on the smaller training set (11,391 sequences) were found to perform marginally better on initial tests, both in terms of the framing and error classification of sequences. Therefore, the nucleotide and amino acid PHMMs trained on 11,391 training sequences were retained for internal use in the coil package. The runtime (on a virtual machine hosted on Compute Canada cloud resources, with 32 CPU cores (2.4GHz) and 120Gb of RAM) for training of the final nucleotide PHMM with 11,391 input sequences was 2 hours and 22 minutes, and the time for training of the corresponding amino acid model was 5 minutes.

### Assessment of coil package performance

Following successful model training and package construction, a random subsample of 10,000 barcode sequences from the previously withheld validation data was used to assess the performance of coil, in terms of its ability to place sequences in the correct reading frame and determine if sequences contain insertion or deletion errors. Additionally, we aimed to test coil’s establishment of reading frame and proper translation when excess sequence information in addition to the COI-5P region is present or when barcode information is missing from the edges of sequences. The 10,000 samples were randomly split and altered using a custom Python script (Supplementary File 2) to construct the following 5 groups (2000 sequences each):

a. No changes made.
b. 1-100 random base pairs added to the front and/or back of the sequence, with equal addition probabilities for each of the four nucleotides.
c. 1-100 base pairs removed from the front and/or back of the sequence.
d. Altered with a random indel probability of 1% per bp (equal probability of either an insertion or deletion, in the case of insertions a random bp was inserted). Point mutations were also applied with a probability of 1% per bp (a 1% chance of a nucleotide being changed to any other nucleotide) in order to produce more realistic error profiles and test indel error detection in the presence of additional noise.
e. Sequences either shortened or elongated in the manner of groups (b) or (c), with errors then introduced in the manner of group (d).

These validation data were then loaded into R and passed through the coil analysis pipeline twice (Supplementary File 3). First, the translation step of the coil pipeline was run using the genetic codes known to correspond to the sequences, and second, the pipeline was run using the default behaviour, assuming unknown genetic codes and utilizing the censored translation option.

The outputs of the pipeline were assessed to determine how well sequences were framed and translated. The package’s ability to frame sequences was evaluated by looking for stop codons in the translation outputs of the 6000 validation sequences with no artificially introduced indel errors. The presence of stop codons would indicate a sequence was not set in reading frame properly (as their original amino acid sequences are known and do not contain stop codons), whereas the absence of stop codons is strongly suggestive of correct translation. Additionally, the log likelihood values obtained from comparison of all 10,000 sequences to the amino acid PHMM were assessed to determine the log likelihood’s effectiveness as an error classifier. Sequences with introduced indel errors and those without errors were labelled in a binary fashion. The R package ROCit (CRAN.R-project.org/package=ROCit) was used to determine the ability of the amino acid PHMM output log likelihood values to discern sequences with errors from those without errors. Comparisons were made between the following groups of validation data: full length sequence data ((a) vs (d)), variable length sequence data ((b) and (c) vs (e)), and all sequence data ((a)(b)(c) vs. (d)(e)). To determine the effect of sequence length on classification ability, the long sequences (b) were compared to the sequences from the variable length error dataset (e) that were elongated, and the short sequences (c) were compared to the sequences from the variable length error dataset (e) that were shortened. The area under the curve (AUC – a metric of how well the log likelihood values separate the two categories) and the optimal F score threshold (the log likelihood value that optimizes the trade-off between false positive and false negative classifications) were determined for all comparisons.

To demonstrate how coil can be applied in metabarcoding analyses, the coil analysis pipeline described above was rerun, and associated evaluation metrics were generated, for non-full-length barcode sequences from a defined subsection for the COI-5P region. Subsets of the barcode sequences in categories (a) and (c) were taken for an arbitrarily chosen 300bp window (base pairs 337-636 of the full COI-5P region) to simulate non-full-length COI-5P sequences commonly generated in DNA metabarcoding. Sequences from group (c) were only retained if they contained a documented indel error within the defined 300bp region (1868 sequences). To determine coil’s baseline classification ability on these data, the coil analysis pipeline was run for the non-full-length sequences using the full-length nucleotide and amino acid PHMMs. To test if the use of targeted PHMMs (corresponding to only that specific 300bp section COI-5P region) improves coil’s ability to frame sequences and identify indels, the complete coil analysis pipeline was then rerun an additional time using a subsection of the PHMMs representing only the given 300bp window to evaluate the non-full-length barcode sequences (Supplementary File 4).

### Processing non-animal barcode sequences

In addition to identifying the presence of indels in animal COI-5P sequences (which would most likely result from technical errors), we tested how coil performed on COI-5P sequences for which it was not designed to process. A series of plant, fungi, and bacterial COI-5P sequences as well as animal nuclear mitochondrial pseudogenes were analyzed with coil, to see if sequences from these categories would be flagged as likely containing errors, or if they would be processed in the same manner as animal COI-5P sequences. The first set of sequences investigated were COI-5P sequences from the plant and fungi kingdoms. The NCBI nucleotide database (https://www.ncbi.nlm.nih.gov/nucleotide/) was queried for ‘cytochrome c oxidase 1’, and the results were filtered to retain sequences from only plants and fungi. Results were filtered by sequence length (minimum 550bp, maximum 2000bp), and the sequence descriptions were all manually checked to ensure that all results were COI-5P sequences. This resulted in a set of 570 plant and 174 fungal sequences (Supplementary File 5) that were then analyzed using the coil analysis pipeline (coi5p_pipe function) with default parameters.

*Wolbachia* is a genus of bacterial endosymbionts (commonly found in arthropods) that can in some instances confound attempts to isolate an animal specimen’s barcode sequence (Smith *et al.* 2012). To see if coil would flag *Wolbachia* sequences as errors (either due to a low likelihood value or the presence of a stop codon in the amino acid sequence), data associated with the accession numbers of *Wolbachia* samples listed in Supplementary File S2 of Smith *et al.* (2012) were obtained from NCBI. Data were filtered to omit sequences <550bp in length, yielding 345 *Wolbachia* sequences that were analyzed via coil’s ‘coi5p_pipe’ function (parameters: trans_table = 0 and indel_threshold = −440.24).

Nuclear mitochondrial pseudogenes (numts) are non-encoded sequences in the nucleus that are the product of mitochondrial DNA fragments being incorporated into the nuclear genome (Bensasson *et al.* 2001). Numts resulting from the COI-5P gene have been shown to interfere with accurate assessment of the COI-5P sequences for a species complex of Australian stingless bees (*Tetragonula carbonaria*, *T. davenporti*, *T. hockingsi*, and *T. mellipes*; hereafter the ‘Carbonaria group’) (Françoso *et al.* 2019). Sequence data for the 9 numts with reported accession numbers in Figure 7 of Françoso *et al.* (2019) were obtained and analyzed with coil (using the coi5p_pipe function and default parameters) to see if the package identified these sequences as likely being erroneous.

**Fig. 7.**
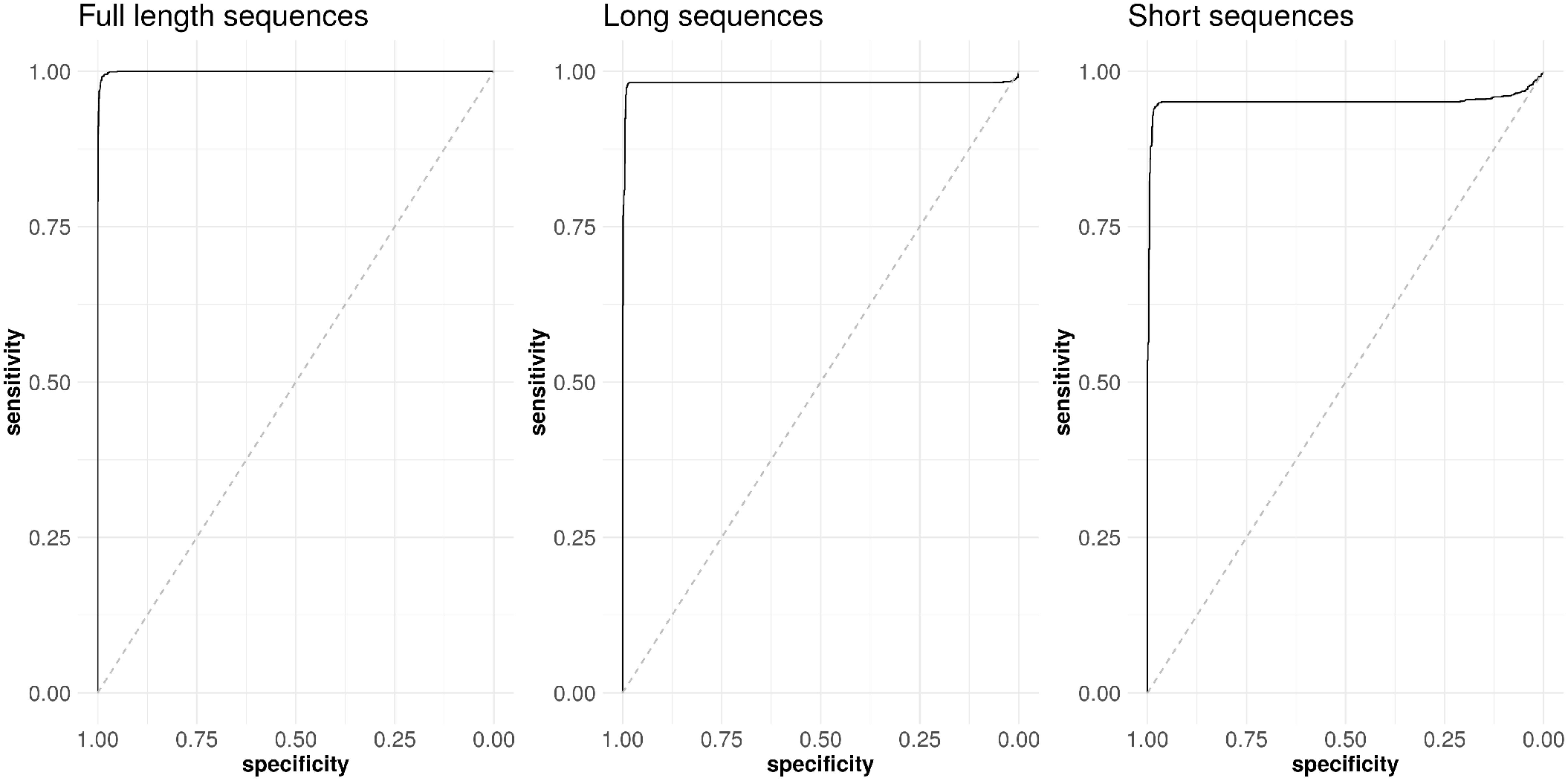
ROC (receiver operating characteristic) curves showing the ability of coil to classify correctly sequences as being correct or containing insertion or deletion errors. Indel classification ability for three types of sequences (all translated with known genetic codes) are displayed: full-length barcode sequences (barcode sequences of unaltered lengths), long sequences (1-100bp added to both ends of the sequences), and short sequences (1-100bp removed from each end of the sequence). Black lines indicate the ability of the classifier to separate true sequences from those with indel errors. A perfect classifier would have an area under the curve (AUC) of 1 and intersect the top left corner of the graph (See Table 1 for corresponding AUC values). The dotted grey lines represent classification through chance (AUC = 0.5).

**Table 1.** Classification accuracy of coil’s indel check function on test sequence groups of different compositions. AUC – area under the curve for the ability of the amino acid PHMM log likelihood score to discriminate sequences with insertion or deletion errors from sequences without errors. The optimal LogL threshold for differentiating sequences with indels from those without indels is shown. The optimal LogL cut off, as determined through F-score maximization, is shown.

| Dataset | Translation | AUC | Optimal LogL Cut Off |
| --- | --- | --- | --- |
| Full length<br>sequences | Genetic code known | 0.9989 | -246.20 |
|  | Censored | 0.9982 | -358.88 |
| Variable<br>length<br>sequences | Genetic code known | 0.9662 | -354.44 |
|  | Censored | 0.9659 | -457.39 |
| Short length<br>sequences | Genetic code known | 0.9509 | -354.44 |
|  | Censored | 0.9504 | -440.24 |
| Long length<br>sequences | Genetic code known | 0.9802 | -324.37 |
|  | Censored | 0.9795 | -412.88 |
| All data<br>combined | Genetic code known | 0.9776 | -308.05 |
|  | Censored | 0.9773 | -425.21 |

## Results & Discussion

### Framing and Translation

The validation data outputs were assessed to determine how effective coil was at processing variable-length barcode data. The package’s ability to set sequences in reading frame was evaluated by looking for stop codons in the translation outputs of the 6000 validation sequences with no introduced indel errors. If stop codons were present, the sequence would not have been set in reading frame properly, whereas the absence of stop codons is strongly suggestive of a correct reading frame establishment. Of the 6000 validation sequences without introduced errors, 5888 (98.13%) were correctly framed. The sequences with incorrectly identified reading frames were not evenly distributed across the three sequence length categories (short sequence, full-length sequences, and long sequences). All barcode sequences in the full-length (a) category except for one were correctly placed in reading frame, indicating that the nucleotide PHMM is able to recognize and frame taxonomically variable barcode data and that it is not biased against certain taxonomic groups. The program appeared to be effective at trimming artificially lengthened sequences (category b) and placing them in reading frame, with only 38 framing errors in the 2000 long sequences (98.1% accuracy). The program was slightly less effective at placing shortened sequences (category c) in reading frame, with 98 errors observed (95.1% accuracy). Despite this imperfect performance, the program is effective at framing the vast majority of sequences and can aid the user by successfully automating this task for most inputs (Supplementary File 6). The summary data provided by the indel check step (stop codon presence and error likelihood values) can aid users in targeting additional correction effort to the subset of sequences coil is unable to place in reading frame.

### Indel Error Identification

Across the entire validation dataset, coil’s indel_check function was able assess the amino acid data and accurately discern sequences with insertion or deletion errors from correct barcode sequences with greater than 97% accuracy. As with the framing of sequences, coil’s effectiveness was influenced by the length of the sequences (Table 1). When the full-length barcode sequences with and without introduced errors were evaluated, they could be categorized with greater than 99.9% accuracy. For sequences with excess DNA sequence added to the ends, the sequences with errors could be discerned from the correct sequences with 98.0% accuracy. For the short sequences, performance was once again the lowest, with 95.1% classification accuracy (Fig. 7). The similarity in these accuracy scores to the percentage of sequences placed in reading frame correctly is not coincidental, but rather due to the fact that the ability to correctly classify a sequence is highly dependent on the ability of coil to place it in the common reading frame correctly.

Closer examination of the classification errors in the short sequence dataset (the category with the highest error frequency) helped to characterize the limitations of coil’s classification ability. The majority of the classification errors (103/132 – 78%) were false negatives, i.e. sequences without introduced errors that were flagged as likely to contain errors. The reason for this appears to be the incorrect establishment of the reading frame and thus large-scale reading frame shifts, resulting in incorrect amino acid sequences. The major limitation of coil therefore appears to be the inability to always establish the reading frame properly, a problem that is magnified in short sequences where the front of the sequence is truncated. The examination of the 29 falsely accepted sequences for the short category revealed the three main patterns where coil failed to detect errors in a sequence: 1) When an indel occurs late (>500 bp) in the sequence (observed for 7 sequences), 2) when three insertions or three deletions are found in closely proximity (observed for 4 sequences), and 3) when an insertion and deletion occur in close proximity to one another (observed for 18 instances). All three of these patterns do not lead to large-scale alterations to the amino acid sequence, and are therefore not easily detected by examination of the amino acid sequence.

In all error classification tests, the accuracy (as shown by AUC) was highly similar when translation was conducted with the known genetic codes and when censored translation was employed (Table 1). This indicates that the censored translation method is effective at allowing error identification to proceed with accuracy in instances where the taxonomic origin of a sequence is unknown. It should be noted that the log likelihood values associated with sequences are overall slightly lower when generated using censored translation, because lower numbers of amino acids contribute to the likelihood value (5/64 codons are not translated – Fig. 5). For this reason, the threshold used to identify sequences with indel errors should be adapted to the characteristics of the data being analyzed. Table 1 provides users with a starting point for selecting a classification threshold, showing the optimal log likelihood threshold (as determined through F-score maximization) for the different comparisons of validation data. These values serve as statistically informed starting points from which a user can deviate if they wish to attempt minimization of type 1 (lowering the logL threshold to minimize rejection of correct sequences) or type 2 error (raising the logL threshold to minimize acceptance of sequences with indel errors).

### Extension of coil to metabarcode data analysis

The indel identification tests showed that coil’s performance is suboptimal when the length of the examined sequence is far shorter than the PHMM profile length. The proximal cause of this issue is that similarity between probabilistic profiles for different parts of the barcode region can lead to continuous series of base pairs from the sequence matching to an incorrect portion of the PHMM and incorrect establishment of the reading frame. This is problematic in the analysis of metabarcode data, where non-full-length (~200-400bp) sections of the COI-5P barcode are targeted for amplification and sequencing by custom primers. The fact that these non-full-length barcodes are in most instances derived from targeted subsections of the COI-5P region means that the framing limitation can be overcome by comparing sequences to only a subsection of coil’s complete PHMMs. Using a subsection of the full PHMMs reduces or eliminates the discrepancy in size between the PHMM and the non-full-length barcodes, drastically reducing the false positive indel signals caused by incorrect establishment of the reading frame.

Using the subsection of the complete PHMMs (defined using coil’s subsetPHMM function), coil was able to separate non-full-length barcode sequences with indel errors from correct sequences with greater than 99.4% accuracy (Fig 8.). This proved to be a drastic improvement over the use of the full-length PHMMs in the analysis of the same 300bp sequence dataset (AUC = 0.77). Using the full-length model, 513 sequences from group (a) were incorrectly flagged as containing stop codons, indicating a high rate of incorrect reading frame establishment (25.6%). This high level of incorrect framing is a result of the 300bp window starting late (base pair 337) in the COI-5P region, providing a large set of leading profile positions to which incorrect matches could occur. Conversely, the false positive rate using the subsection of the full PHMMs led to an extremely low false positive rate (0.1%). This indicates that the coil analysis pipeline is appropriate for technical error identification in non-full-length metabarcode outputs, if there is available information on the approximate subsection of the COI-5P region from which the sequences are derived.

**Fig 8.**
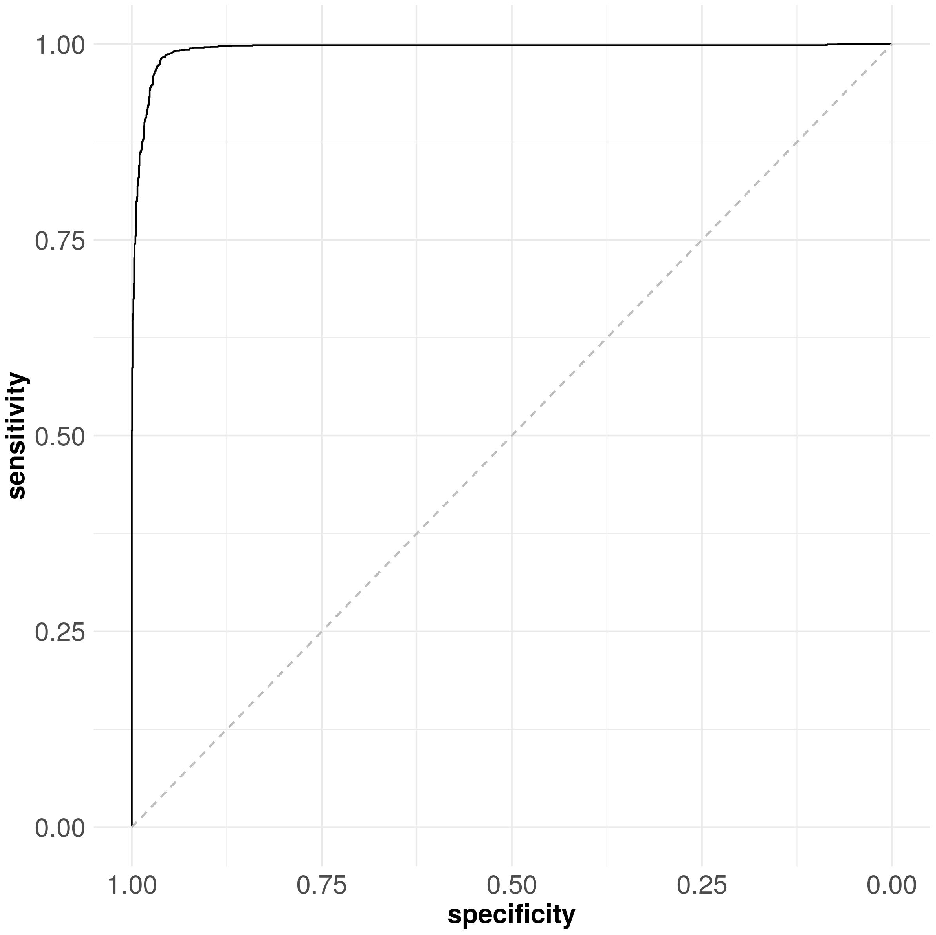
ROC (receiver operating characteristic) curve showing coil’s ability to classify correctly non-full-length barcode sequences (300bp) from a defined sub-region of the COI-5P barcode, using a subsection of the nucleotide and amino acid PHMMs corresponding to the same sub-region. The ROC values shown correspond to analysis using censored translation, which assumes unknown genetic code (AUC = 0.994). When the analysis was repeated using known genetic codes, the AUC value improved slightly (AUC = 0.996).

It should also be noted that amino acid PHMM log likelihood values reported for input sequences are dependent on the size of the model to which they are compared. The optimal log likelihood cut offs presented (in Table 1) are therefore not applicable when user-defined subsets of the complete PHMMs are used. Supplementary File 4 has been included to aid in determining the optimal log likelihood cut off for a user defined COI-5P subsection of interest.

### Package limitations

The results of processing non-animal COI-5P sequences with coil were variable; 167/174 (96%) of fungi and 336/570 (58.8%) of plant sequences were flagged as likely erroneous based on the resulting log likelihood scores and/or the presence of stop codons. The reason for coil labelling fungi sequences as likely erroneous is the existence of introns in fungal COI-5P sequences (Avin *et al.* 2017). Fungal sequences show drastic variability in length due to the presence of introns, compared to animal sequences which do not contain introns and are relatively consistent in length (Avin *et al.* 2017). This variation in the structure of fungal sequences is interpreted by coil’s processing pipeline as evidence of indel errors, resulting in the majority of true fungal sequences being flagged as erroneous. Plant COI-5P sequences are less variable in structure than fungi, but more than half of the sequences tested deviated from the animal COI-5P profile in a manner extreme enough for them being flagged as erroneous. These results show that coil cannot presently be used to pre-process and filter errors reliably from non-animal COI-5P sequences. The PHMMs are not trained to accommodate non-animal data, and processing sequences from the plant or fungal kingdoms results in a large number of false negatives. A future extension of coil can be the addition of PHMMs trained on data from different kingdoms or barcodes, which would allow coil’s processing pipeline to be applied in error filtration of other protein-coding, intron-free barcodes.

Of the 345 *Wolbachia* COI-5P sequences analyzed, only five were flagged as likely to be erroneous, and only a single sequence contained stop codons. The five instances of *Wolbachia* sequences being flagged as errors resulted from the incorrect framing and translation of sequences. Since the *Wolbachia* sequences are real instances of the COI-5P gene (albeit from bacteria as opposed to animals) that do not possess any drastic changes to reading frame relative to animals, their nucleotide and corresponding amino acid sequences are likely enough to be accepted as matches to the animal COI-5P PHMMs. This demonstrates that coil cannot be effectively appropriated to separate animal COI-5P sequences from the COI-5P sequences of endosymbiote contaminants.

All 9 of the numt sequences from the Carbonaria group analyzed with coil were flagged as likely to contain an error. This was accomplished despite the fact that none of the translated sequences contained a stop codon. Each pseudogene possessed an amino acid sequence that the program characterized as improbable (all log likelihood values lower than −754.00). It should be noted that the numts analyzed were shorter sequences (333bp each), which inherently possess lower log likelihood values. However, animal COI-5P sequence fragments of this length tend to yield log likelihood values of approximately −450.00, so the drastically lower log likelihood sequences are sufficient to distinguish the pseudogene sequences from COI-5P sequences. The number of numts tested was low and limited to sequences identified within a single species complex. Expansion of these tests to COI-5P-derived numts from a more diverse set of taxa could reveal how well coil is able to characterize numts. At present, it is hypothesized that the ability to discern pseudogenes from true barcode sequences will depend on how diverged the structure and sequence of a pseudogene are, with frame-shift-causing indels increasing the likelihood of detection by coil.

Taken together, these results show that coil Version 1.0 should only be used to process animal COI-5P data and that it is less reliable at detecting biological contaminants than technical errors. The genetic structure of biological contaminants such as *Wolbachia* COI-5P is not always dissimilar enough from animal COI-5P sequences for the PHMMs to discern the two categories. Additionally, if an animal specimen were contaminated with DNA not from another kingdom, but rather with DNA from another animal, then detection with coil in most instances would not be possible, as the models employed by the package are trained on a taxonomically robust sample of animal sequences. The results presented here regarding the identification of biological contaminants are conservative, as in all tests we have used coil’s default censored translation option, which assumes that taxonomic information is not available. The ability to identify biological contaminants can be greatly increased if the taxonomy (and therefore the genetic code) corresponding to a sample is known. Contaminants from organisms that appropriate a different genetic code would have a higher likelihood of being identified in this situation, as translation would be more likely to yield amino acid sequences with stop codons or highly improbable amino acids. Overall the coil package should not be considered a panacea for all DNA barcode data errors, but rather as a tool that can be incorporated into barcode analysis as a reliable pre-processing and data-cleaning step that can identify technical errors. Researchers using barcode data should consider additional steps in their analyses, such as BLAST, which can help inform the taxonomy of sequences and identify biological contaminants.

## Conclusion

We have presented the R package coil and demonstrated how it can effectively set COI-5P barcode data in reading frame, translate sequences in a taxonomically robust fashion, and identify sequences containing frame-shifting indel errors with greater than 97% accuracy. A vignette demonstrating the software’s functionality and suggested workflows is available within the package (and included here as well – Supplementary File 7). The package can facilitate exploratory analysis of barcode data and aid in data cleaning and error assessment. At present, the package’s functionality is limited to the main DNA barcode region for animals, COI-5P. In the future, we hope to extend the package’s functionality to additional DNA barcode regions, including alternative animal barcodes like cytochrome b (Kochzius *et al.* 2010; Yacoub *et al.* 2015), as well barcodes employed in analysis of species from other kingdoms (Hollingsworth *et al.* 2009; Hollingsworth *et al.* 2011). Adding this functionality would require the training of addition profile hidden Markov models for these barcode regions and integration of these models into the coil package. This is a feasible addition thanks to the large amount of additional barcode data available through BOLD (Ratnasingham & Hebert 2007) and other protein-coding markers available through NCBI, which can be queried and filtered using methods similar to those employed here for COI-5P. Another extension of the functionality of the coil package is to move from error identification to error correction. PHMMs are robust models of the conserved sequence of the COI-5P barcode region on both a DNA and amino acid level. Accurately identifying where deviations from the expected sequence occur and applying corrections to the sequences in an unbiased fashion would allow PHMMs to serve as the basis of a DNA barcode data denoiser. Corrections based on PHMMs trained on the wealth of barcode sequence information available could serve as a powerful complement, or alternative, to current denoising methods which rely on read abundance and *de novo* clustering of sequences (Nearing *et al.* 2018).

## Acknowledgements

Funding for this research was provided by grants in Bioinformatics and Computational Biology from the Government of Canada through Genome Canada and Ontario Genomics and from the Ontario Research Fund. Funders played no role in software design or manuscript preparation. This research was enabled in part by resources provided by Compute Canada (www.computecanada.ca). Thank you to Samantha E. Majoros for aiding in the initial testing of the coil package.

## List of Supplementary Files

**Supplementary File 1** – ‘S1-train_bold_coi_public_data_subsample.zip’ A .tsv file with the representative subsample of the BOLD public database used to train the nucleotide and amino acid PHMMs.

**Supplementary File 2** – ‘S2-create_assessment_data.py’ Custom python script used in the creation of the variable quality dataset for testing coil.

**Supplementary File 3** – ‘S3-coil-utilization.r’ R script for application of the coil pipeline in the analysis of the validation data.

**Supplementary File 4** – ‘S4-constrained_eval.r’ R script that contains the code for analyzing short 300bp barcode sequences with subsets of the complete PHMMs.

**Supplementary File 5** – ‘S5-non_animal_tests_output.tsv’ Output from the analysis of publicly available *Wolbachia,* plant and fungi COI-5P barcode sequences and nuclear mitochondrial pseudogenes (numts) using the coil analysis pipeline.

**Supplementary File 6** – ‘S6-validation_data’ Folder containing the validation data inputs and outputs produced for the assessment of coil effectiveness.

**Supplementary File 7** – ‘S7-coil-vignette.pdf’ A pdf demonstrating the functionality of the coil package. This is the static version of the package vignette, which is available through R. The original R markdown file is available for download from: https://github.com/CNuge/coil/tree/master/vignettes

## References

Avin, F.A., Subha B., Tan Y.S., Braukmann T.W., Vikineswary S., Hebert P.D.N. 2017. Escaping introns in COI through cDNA barcoding of mushrooms: *Pleurotus* as a test case. Ecology and Evolution. 7(17):6972–80. doi:10.1002/ece3.3049.

Bensasson D., Zhang D.X., Hartl D.L., Hewitt G.M. 2001. Mitochondrial pseudogenes: evolution’s misplaced witnesses. Trends in Ecology – Evolution. 16(6):314–21. doi:10.1016/s0169-5347(01)02151-6.

Calderón-Sanou I., Münkemüller T., Boyer F., Zinger L., Thuiller W. 2019. From environmental DNA sequences to ecological conclusions: How strong is the influence of methodological choices?. Journal of Biogeography. 47(1):193–206. doi:10.1111/jbi.13681.

Castresana J., Lübben M., Saraste M., Higgins D.G. 1994. Evolution of cytochrome oxidase, an enzyme older than atmospheric oxygen. The EMBO journal. 13(11):2516–25. doi:10.1002/j.1460-2075.1994.tb06541.x

Charif D., Lobry J.R. 2007. SeqinR 1.0-2: a contributed package to the R project for statistical computing devoted to biological sequences retrieval and analysis. In Structural approaches to sequence evolution 2007 (pp. 207–232). Springer, Berlin, Heidelberg. doi:10.1007/978-3-540-35306-5_10

Cristescu M.E. 2014. From barcoding single individuals to metabarcoding biological communities: towards an integrative approach to the study of global biodiversity. Trends in Ecology & Evolution. 29(10):566–71. doi:10.1016/j.tree.2014.08.001

Durbin R., Eddy S.R., Krogh A., Mitchison G. 1998. Biological sequence analysis: probabilistic models of proteins and nucleic acids. Cambridge University Press, Cambridge.

Eddy S.R. 2009. A new generation of homology search tools based on probabilistic inference. In Genome Informatics Series. Vol. 23 pp. 205–211.

Edgar R.C. 2010. Search and clustering orders of magnitude faster than BLAST. Bioinformatics. 26(19):2460–1. doi:10.1093/bioinformatics/btq461.

Elbrecht V., Vamos E.E., Steinke D., Leese F. 2018. Estimating intraspecific genetic diversity from community DNA metabarcoding data. PeerJ. 6:e4644. doi:10.7717/peerj.4644.

Elzanowski A., Ostell J. 2019. The Genetic Codes [online]. Available from https://www.ncbi.nlm.nih.gov/Taxonomy/Utils/wprintgc.cgi#SG5. [accessed Sept 9 2019].

Françoso E., Zuntini A.R., Ricardo P.C., Silva J.P., Brito R., Oldroyd B.P., Arias M.C. 2019. Conserved numts mask a highly divergent mitochondrial-COI gene in a species complex of Australian stingless bees *Tetragonula* (Hymenoptera: Apidae). Mitochondrial DNA Part A. 16:1–2. doi:10.1080/24701394.2019.1665036.

Folmer O., Black M., Hoeh W., Lutz R., Vrijenhoek R. 1994. DNA primers for amplification of mitochondrial cytochrome c oxidase subunit I from diverse metazoan invertebrates. Molecular Marine Biology and Biotechnology. 3(5):294–9. PMID: 7881515.

Hebert P.D.N., Cywinska A., Ball S.L., deWaard J.R. 2003. Biological identifications through DNA barcodes. Proceedings of the Royal Society of London. Series B: Biological Sciences. 270(1512):313–21. doi:10.1098/rspb.2002.2218.

Hebert P.D.N., Penton E.H., Burns J.M., Janzen D.H., Hallwachs W. 2004. Ten species in one: DNA barcoding reveals cryptic species in the neotropical skipper butterfly *Astraptes fulgerator*. Proceedings of the National Academy of Sciences. 101(41):14812–7. doi:10.1073/pnas.0406166101.

Hebert P.D.N., Braukmann T.W., Prosser S.W., Ratnasingham S., deWaard J.R., Ivanova N.V., Janzen D.H., Hallwachs W., Naik S., Sones J.E., Zakharov E.V. 2018. A Sequel to Sanger: amplicon sequencing that scales. BMC Genomics. 19(1):219. doi:10.1186/s12864-018-4611-3.

Hollingsworth P.M., Forrest L.L., Spouge J.L., Hajibabaei M., Ratnasingham S., van der Bank M., Chase M.W., Cowan R.S., Erickson D.L., Fazekas A.J. 2009. A DNA barcode for land plants. Proceedings of the National Academy of Sciences. 106(31):12794–7. doi:10.1073/pnas.0905845106.

Hollingsworth P.M., Graham S.W., Little D.P. 2011. Choosing and using a plant DNA barcode. PloS One. 6(5):e19254. doi:10.1371/journal.pone.0019254.

Hubert N., Hanner R. 2015. DNA barcoding, species delineation and taxonomy: a historical perspective. DNA Barcodes. 3(1):44–58. doi:10.1515/dna-2015-0006.

Jukes T.H., Osawa S. 1993. Evolutionary changes in the genetic code. Comparative Biochemistry and Physiology. B, Comparative Biochemistry. 106(3):489–94. doi:10.1016/0305-0491(93)90122-l.

Kochzius M., Seidel C., Antoniou A., Botla S.K., Campo D., Cariani A., Vazquez E.G., Hauschild J., Hervet C., Hjörleifsdottir S., Hreggvidsson G. 2010. Identifying fishes through DNA barcodes and microarrays. PLoS One. 5(9):e12620. doi:10.1371/journal.pone.0012620

Nearing J.T., Douglas G.M., Comeau A.M., Langille M.G. 2018. Denoising the Denoisers: an independent evaluation of microbiome sequence error-correction approaches. PeerJ. 6:e5364. doi:10.7717/peerj.5364.

Osawa S., Jukes T.H., Watanabe K., Muto A. 1992. Recent evidence for evolution of the genetic code. Microbiology and Molecular Biology Reviews. 56(1):229–64. PMCID: PMC372862.

Paradis E., Schliep K. 2018. ape 5.0: an environment for modern phylogenetics and evolutionary analyses in R. Bioinformatics. 35(3):526–8. doi:10.1093/bioinformatics/bty633.

Pentinsaari M., Salmela H., Mutanen M., Roslin T. 2016. Molecular evolution of a widely-adopted taxonomic marker (COI) across the animal tree of life. Scientific Reports. 13;6:35275. doi:10.1038/srep35275.

Porter T.M., Hajibabaei M. 2018. Over 2.5 million COI sequences in GenBank and growing. PLoS One. 13(9):e0200177. doi:10.1371/journal.pone.0200177.

Ranwez V., Harispe S., Delsuc F., Douzery E.J. 2011. MACSE: Multiple Alignment of Coding SE-quences accounting for frameshifts and stop codons. PLoS One. 6(9):e22594. doi:10.1371/journal.pone.0022594.

Ratnasingham S., Hebert P.D.N. 2007. BOLD: The Barcode of Life Data System (http://www.barcodinglife.org). Molecular Ecology Notes. 7(3):355–64. doi:10.1111/j.1471-8286.2007.01678.

Schirmer M., D’Amore R., Ijaz U.Z., Hall N., Quince C. 2016. Illumina error profiles: resolving fine-scale variation in metagenomic sequencing data. BMC Bioinformatics. 17(1):125. doi:10.1186/s12859-016-0976-y

Schoch C.L., Seifert K.A., Huhndorf S., Robert V., Spouge J.L., Levesque C.A., Chen W., Fungal Bar-coding Consortium. 2012. Nuclear ribosomal internal transcribed spacer (ITS) region as a universal DNA barcode marker for Fungi. Proceedings of the National Academy of Sciences. 109(16):6241–6. doi:10.1073/pnas.1117018109.

Stoeckle M.Y, Kerr K.C. 2012. Frequency matrix approach demonstrates high sequence quality in avian BARCODEs and highlights cryptic pseudogenes. PLoS One. 7(8). doi:10.1371/journal.pone.0043992

Tsukihara T., Aoyama H., Yamashita E., Tomizaki T., Yamaguchi H., Shinzawa-Itoh K., Nakashima R., Yaono R., Yoshikawa S. 1995. Structures of metal sites of oxidized bovine heart cytochrome c oxidase at 2.8 A. Science. 269(5227):1069–74. doi:10.1126/science.7652554

Wilkinson S.P. 2019. aphid: an R package for analysis with profile hidden Markov models. Bioinformatics. 35(19):3829–30. doi:10.1093/bioinformatics/btz159.

Yacoub H.A., Fathi M.M., Sadek M.A. 2015. Using cytochrome b gene of mtDNA as a DNA barcoding marker in chicken strains. Mitochondrial DNA. 26(2):217–23. doi:10.3109/19401736.2013.825771

Youle R.J. 2019. Mitochondria—Striking a balance between host and endosymbiont. Science. 365(6454):eaaw9855. doi:10.1126/science.aaw9855.

