## supplementary for "coil: an R package for cytochrome C oxidase I (COI) DNA barcode data cleaning, translation, and error evaluation": S2-create_assessment_data.py

```
import sys
import numpy as np
import pandas as pd
import random
from numpy.random import random_sample, randint

def random_add(seq_str , side = 0):
	"""
	remove betweem 1 and 100 bp from sequence end.
	trim side: 0 means both, 1 means front and 2 means back.
	"""
	dna = ["a","g","c","t"]

	if side == 0:
		#add both
		front = ''.join([random.choice(dna) for x in range(random.randint(0,100))])
		back = ''.join([random.choice(dna) for x in range(random.randint(0,100))])
		seq_str = front + seq_str + back

	elif side == 1:
		#add front
		front = ''.join([random.choice(dna) for x in range(random.randint(0,100))])
		seq_str = front + seq_str 

	elif side == 2:	
		#add back
		back = ''.join([random.choice(dna) for x in range(random.randint(0,100))])
		seq_str = seq_str + back

	return seq_str


def random_trim(seq_str , side = 0):
	"""
	Remove betweem 1 and 100 bp from sequence end.
	trim side: 0 means both, 1 means front and 2 means back.
	"""
	if side == 0:
		#trim both
		seq_str = seq_str[random.randint(0,100):(len(seq_str) - random.randint(0,100))]		

	elif side == 1:
		#trim front
		seq_str = seq_str[random.randint(0,100):]		

	elif side == 2:	
		#trim back
		seq_str = seq_str[:(len(seq_str) - random.randint(0,100))]		

	return seq_str


def random_bp():
	""" 
	Returns a random nucleotide.
	"""
	bp = ['a', 't', 'g', 'c']
	return random.choice(bp)


def random_mutation(current):
	""" 
	Returns a random nucleotide that is not the same as the current nucleotide.
	"""
	all_bp = ['a', 't', 'g', 'c']
	bp = [b for b in all_bp if b != current]
	return random.choice(bp)


def enumerate_strings(dna_string):
	"""
	Take a string of DNA and compress the homopolymers into counts.
	input: 'ATTTCGGC'
	output:[('A',1), ('T',3), ('C', 1), ('G',2), ('C',1)]
	"""
	seq_dat = list(dna_string)

	#initiate the structure with the first base pair in the sequence
	compressed_list = [[seq_dat[0], 1]]

	for x in seq_dat[1:]:
		if x == compressed_list[-1][0]:
			compressed_list[-1][1] += 1
		else:
			compressed_list.append([x, 1])			

	return compressed_list


def error_introduce(dna_string, global_mutation_rate = 0.01, global_indel_rate = 0.01):
	"""
	Introduce mutations and indel errors into the DNA string.
	"""
	new_seq = ''
	inserton_positions = []
	deletion_positions = []

	enum_seq = enumerate_strings(dna_string)
	org_position = 1 #will record with a 1 index, as the alignment processing will be in r
	# iterate_through the compressed positions in the sequence
	for dat in enum_seq:
		bp = dat[0]
		count = dat[1]	
		#treat each base pair with the overall error profile
		for base in range(count):
			org_position += 1
			val = random_sample()
			#if random number less than threshold 1, do a mutation
			if val < global_mutation_rate:
				new_seq += random_mutation(bp)
			#if the random number is between thresholds 1 and 2, then do an indel
			elif val < (global_indel_rate + global_mutation_rate):
				#0 = insert, 1 = delete
				mut_type = randint(2)
				if mut_type == 0:
					new_seq += bp
					new_seq += random_bp()
					inserton_positions.append(org_position)
				else:
					#add nothing to seq, we skip this bp
					deletion_positions.append(org_position)
			#just add the normal base
			else:
				new_seq += bp

	return (new_seq , 
			';'.join([str(x) for x in inserton_positions]), 
			';'.join([str(x) for x in deletion_positions]))


def trim_or_add_and_mutate(seq, change = 0):
	"""
	Either trim or add sequence prior to introducing errors.
	0 =  add
	1 = delete
	"""
	if change == 0:
		seq = random_add(seq , random.randint(0,2))
	elif change == 1:
		seq = random_trim(seq , random.randint(0,2))
	seq = error_introduce(seq)

	return seq

if __name__ == '__main__':
	"""
	original big raw validation data file was imported here and randomly subset.
	for a demonstration of running this file you can load:
	val_dat = pd.read_csv('S4-validation_data/inputs/sequence_no_changes_2k.csv')
	then applying the functions to val_dat as opposed to mixed_df (etc)
	can demonstrate the execution.

	the functions in this script are generic and can be applied any DNA sequence.
	"""
	validation_file = 'coi5p_validation.tsv'

	val_dat = pd.read_csv(validation_file, sep='\t')

	use_dat = val_dat.sample(10000)

	use_dat['genbankid'][use_dat['genbankid']==11] = 13
	
	len(use_dat['family'].unique()) #1153 families

	keep = ['processid','clean_nucaln',
			'genbankid', 'phylum', 'class',
			'order', 'family', 'genus', 'species',]

	######################################################################
	#2k left alone
	leave_df = use_dat[:2000]

	leave_df = leave_df[keep]

	leave_df['test_seq'] = leave_df['clean_nucaln']

	leave_df.to_csv('data/sequence_no_changes_2k.csv')

	######################################################################
	#2k delete info on edges
	del_df = use_dat[2000:4000]

	del_df = del_df[keep]

	del_df['test_seq'] = del_df.apply(lambda x:
		random_trim(x['clean_nucaln'], random.randint(0,2))
		, axis = 1)

	del_df.to_csv('data/sequence_deletions_2k.csv')

	######################################################################
	#2k add info on edges
	add_df = use_dat[4000:6000]

	add_df = add_df[keep]

	add_df['test_seq'] = add_df.apply(lambda x:
		random_add(x['clean_nucaln'], random.randint(0,2))
		, axis = 1)

	#add_df.test_seq.apply(lambda x: len(x))

	add_df.to_csv('data/sequence_added_2k.csv')

	######################################################################
	#2k indels
	indel_df = use_dat[6000:8000]

	indel_df = indel_df[keep]

	indel_df['error_sequences'] = indel_df.apply(lambda x:
		error_introduce(x['clean_nucaln'])
		, axis = 1)

	indel_df['test_seq'] = indel_df.apply(lambda x:
		x['error_sequences'][0], axis = 1)

	indel_df['inserton_positions'] = indel_df.apply(lambda x:
		x['error_sequences'][1], axis = 1)

	indel_df['deletion_positions'] = indel_df.apply(lambda x:
		x['error_sequences'][2], axis = 1)

	indel_df = indel_df.drop(['error_sequences'], axis=1)

	indel_df.to_csv('data/sequence_indel_2k.csv')

	######################################################################
	#2k add or remove info and have errors.
	mixed_df = use_dat[8000:]

	mixed_df = mixed_df[keep]

	mixed_df['error_sequences']= mixed_df.apply(lambda x:
		trim_or_add_and_mutate(x['clean_nucaln'], random.randint(0,1))
		, axis = 1)

	mixed_df['test_seq'] = mixed_df.apply(lambda x:
		x['error_sequences'][0], axis = 1)

	mixed_df['inserton_positions'] = mixed_df.apply(lambda x:
		x['error_sequences'][1], axis = 1)

	mixed_df['deletion_positions'] = mixed_df.apply(lambda x:
		x['error_sequences'][2], axis = 1)

	mixed_df = mixed_df.drop(['error_sequences'], axis=1)

	mixed_df.to_csv('data/sequence_messy_indel_2k.csv')
```
