## supplementary for "coil: an R package for cytochrome C oxidase I (COI) DNA barcode data cleaning, translation, and error evaluation": S3-coil-utilization.r

```
#install.packages('coil')
library(coil) #manuscript was prepared using version coil 1.0
#install.packages('tidyverse')
library(tidyverse)
library(parallel) #this is a base R package, no need to install

#unzip Supplementary folder #6, and make that the working directory
#Unix command:
#tar -zxvf S6-validation_data.tar.gz
setwd('S6-validation_data')

clean_seq_file = 'inputs/sequence_no_changes_2k.csv'
short_seq_file = 'inputs/sequence_deletions_2k.csv'
long_seq_file = 'inputs/sequence_added_2k.csv'
indel_seq_file = 'inputs/sequence_indel_2k.csv'
messy_seq_file = 'inputs/sequence_messy_indel_2k.csv'

# This detects the number of cores on your machine. 
ncores = detectCores()
# coil is optimally deployed in a parallel fashion (demonstrated below with the mclapply function).
# Testing has shown that processing with coil takes an average of approximately 0.4-0.5 seconds for sequences
# shorter than the full COI-5P region, and 0.8-1 core second per sequence for sequences longer than the full
# COI-5P region. Processing time can be roughly calculated as ((number of sequences/number of cores)/2).
# or in the case of long sequences ((number of sequences/number of cores)). This calculation in presented
# roughly, as it will depend on numerous variables such as the exact specifications of the hardware in question.

# The approximate processing time for a 2K sequence group in the validation data (on a virtual machine hosted on 
# Compute Canada cloud resources, with 32 CPU cores (2.4GHz) and 120Gb of RAM) 
# using iterative execution was 880 - 1600 seconds. Processing of sequences in parallel (using mclapply) across
# the 32 available cores led to processing times between 30 seconds and 1 minute.


###########################################
# clean, known translation tables
###########################################
clean_dat = read_csv(clean_seq_file)

clean_known_translation_data_output = mclapply(1:length(clean_dat$processid), function(i){
   coi5p_pipe(clean_dat$test_seq[i],
              name = clean_dat$processid[i],
              trans_table = clean_dat$genbankid[i])
 },mc.cores = ncores)

clean_known_translation_df = flatten_coi5p(clean_known_translation_data_output)


#Alternative workflow for low memory machine:
# clean_known_translation_df = data.frame(matrix(ncol = 7, nrow = 0),stringsAsFactors = FALSE )
# colnames(clean_known_translation_df) = c("name", "raw", "framed", "aaSeq", "aaScore", "indel_likely", "stop_codons")
#
# progress_bar = txtProgressBar(min=0, max=2000, style = 1, char="=")
# for(i in 1:length(clean_dat$processid)){
# 	out_data = coi5p_pipe(clean_dat$test_seq[i],
# 							name = clean_dat$processid[i],
# 							trans_table = clean_dat$genbankid[i])
# 	clean_known_translation_df = rbind(clean_known_translation_df, flatten_coi5p(list(out_data)))
# 	setTxtProgressBar(progress_bar, value = i)
# }
# close(progress_bar)

write_csv(clean_known_translation_df, "clean_data_known_translation_output.csv")

###########################################
# clean, censored translation
###########################################

clean_censored_translation_data_output = mclapply(1:length(clean_dat$processid), function(i){
  coi5p_pipe(clean_dat$test_seq[i],
             name = clean_dat$processid[i])
},mc.cores = ncores)

clean_censored_translation_df = flatten_coi5p(clean_censored_translation_data_output)

#Alternative workflow for low memory machine:
# clean_censored_translation_df = data.frame(matrix(ncol = 7, nrow = 0),stringsAsFactors = FALSE )
# colnames(clean_censored_translation_df) = c("name", "raw", "framed", "aaSeq", "aaScore", "indel_likely", "stop_codons")
#
# progress_bar = txtProgressBar(min=0, max=2000, style = 1, char="=")
#
# for(i in 1:length(clean_dat$processid)){
# 	out_data = coi5p_pipe(clean_dat$test_seq[i],
# 							name = clean_dat$processid[i])
# 	clean_censored_translation_df = rbind(clean_censored_translation_df, flatten_coi5p(list(out_data)))
# 	setTxtProgressBar(progress_bar, value = i)
# }
# close(progress_bar)

write_csv(clean_censored_translation_df, "clean_data_censored_translation_output.csv")

###########################################
# short, known translation tables
###########################################
short_dat = read_csv(short_seq_file)

short_known_translation_data_output = mclapply(1:length(short_dat$processid), function(i){
  coi5p_pipe(short_dat$test_seq[i],
             name = short_dat$processid[i],
             trans_table = short_dat$genbankid[i])
},mc.cores = ncores)

short_known_translation_df = flatten_coi5p(short_known_translation_data_output)

#Alternative workflow for low memory machine:
# short_known_translation_df = data.frame(matrix(ncol = 7, nrow = 0),stringsAsFactors = FALSE )
# colnames(short_known_translation_df) = c("name", "raw", "framed", "aaSeq", "aaScore", "indel_likely", "stop_codons")
#
# progress_bar = txtProgressBar(min=0, max=2000, style = 1, char="=")
# for(i in 1:length(short_dat$processid)){
# 	out_data = coi5p_pipe(short_dat$test_seq[i],
# 							name = short_dat$processid[i],
# 							trans_table = short_dat$genbankid[i])
# 	short_known_translation_df = rbind(short_known_translation_df, flatten_coi5p(list(out_data)))
# 	setTxtProgressBar(progress_bar, value = i)
# }
# close(progress_bar)

write_csv(short_known_translation_df, "short_data_known_translation_output.csv")

###########################################
# short, censored translation
###########################################
short_censored_translation_data_output = mclapply(1:length(short_dat$processid), function(i){
  coi5p_pipe(short_dat$test_seq[i],
             name = short_dat$processid[i])
},mc.cores = ncores)

short_censored_translation_df = flatten_coi5p(short_censored_translation_data_output)

#Alternative workflow for low memory machine:
# short_censored_translation_df = data.frame(matrix(ncol = 7, nrow = 0),stringsAsFactors = FALSE )
# colnames(short_censored_translation_df) = c("name", "raw", "framed", "aaSeq", "aaScore", "indel_likely", "stop_codons")
#
# progress_bar = txtProgressBar(min=0, max=2000, style = 1, char="=")
# for(i in 1:length(short_dat$processid)){
# 	out_data = coi5p_pipe(short_dat$test_seq[i],
# 							name = short_dat$processid[i])
# 	short_censored_translation_df = rbind(short_censored_translation_df, flatten_coi5p(list(out_data)))
# 	setTxtProgressBar(progress_bar, value = i)
# }
# close(progress_bar)

write_csv(short_censored_translation_df, "short_data_censored_translation_output.csv")

###########################################
# long, known translation tables
###########################################
long_dat = read_csv(long_seq_file)

long_known_translation_data_output = mclapply(1:length(long_dat$processid), function(i){
  coi5p_pipe(long_dat$test_seq[i],
             name = long_dat$processid[i],
             trans_table = long_dat$genbankid[i])
},mc.cores = ncores)

long_known_translation_df = flatten_coi5p(long_known_translation_data_output)

#Alternative workflow for low memory machine:
# long_known_translation_df = data.frame(matrix(ncol = 7, nrow = 0),stringsAsFactors = FALSE )
# colnames(long_known_translation_df) = c("name", "raw", "framed", "aaSeq", "aaScore", "indel_likely", "stop_codons")
#
# progress_bar = txtProgressBar(min=0, max=2000, style = 1, char="=")
# for(i in 1:length(long_dat$processid)){
# 	out_data = coi5p_pipe(long_dat$test_seq[i],
# 							name = long_dat$processid[i],
# 							trans_table = long_dat$genbankid[i])
#
# 	long_known_translation_df = rbind(long_known_translation_df, flatten_coi5p(list(out_data)))
# 	setTxtProgressBar(progress_bar, value = i)
# }
# close(progress_bar)

write_csv(long_known_translation_df, "long_data_known_translation_output.csv")

###########################################
# long, censored translation
###########################################

long_censored_translation_data_output = mclapply(1:length(long_dat$processid), function(i){
  coi5p_pipe(long_dat$test_seq[i],
             name = long_dat$processid[i])
},mc.cores = ncores)

long_censored_translation_df = flatten_coi5p(long_censored_translation_data_output)

#Alternative workflow for low memory machine:
# long_censored_translation_df = data.frame(matrix(ncol = 7, nrow = 0),stringsAsFactors = FALSE )
# colnames(long_censored_translation_df) = c("name", "raw", "framed", "aaSeq", "aaScore", "indel_likely", "stop_codons")
#
# progress_bar = txtProgressBar(min=0, max=2000, style = 1, char="=")
# for(i in 1:length(long_dat$processid)){
# 	out_data = coi5p_pipe(long_dat$test_seq[i],
# 							name = long_dat$processid[i])
#
# 	long_censored_translation_df = rbind(long_censored_translation_df, flatten_coi5p(list(out_data)))
# 	setTxtProgressBar(progress_bar, value = i)
# }
# close(progress_bar)

write_csv(long_censored_translation_df, "long_data_censored_translation_output.csv")

###########################################
# indel, known translation tables
###########################################
indel_dat = read_csv(indel_seq_file)

indel_known_translation_data_output = mclapply(1:length(indel_dat$processid), function(i){
 coi5p_pipe(indel_dat$test_seq[i],
            name = indel_dat$processid[i],
            trans_table = indel_dat$genbankid[i])
},mc.cores = ncores)

indel_known_translation_df = flatten_coi5p(indel_known_translation_data_output)


#Alternative workflow for low memory machine:
# indel_known_translation_df = data.frame(matrix(ncol = 7, nrow = 0),stringsAsFactors = FALSE )
# colnames(indel_known_translation_df) = c("name", "raw", "framed", "aaSeq", "aaScore", "indel_likely", "stop_codons")
#
# progress_bar = txtProgressBar(min=0, max=2000, style = 1, char="=")
#
# for(i in 1:length(indel_dat$processid)){
# 	out_data = coi5p_pipe(indel_dat$test_seq[i],
# 							name = indel_dat$processid[i],
# 							trans_table = indel_dat$genbankid[i])
#
# 	indel_known_translation_df = rbind(indel_known_translation_df, flatten_coi5p(list(out_data)))
# 	setTxtProgressBar(progress_bar, value = i)
# }
# close(progress_bar)

write_csv(indel_known_translation_df, "indel_data_known_translation_output.csv")

###########################################
# indel, censored translation
###########################################

indel_censored_translation_data_output = mclapply(1:length(indel_dat$processid), function(i){
 coi5p_pipe(indel_dat$test_seq[i],
            name = indel_dat$processid[i])
},mc.cores = ncores)

indel_censored_translation_df = flatten_coi5p(indel_censored_translation_data_output)


#Alternative workflow for low memory machine:
# indel_censored_translation_df = data.frame(matrix(ncol = 7, nrow = 0),stringsAsFactors = FALSE )
# colnames(indel_censored_translation_df) = c("name", "raw", "framed", "aaSeq", "aaScore", "indel_likely", "stop_codons")
#
# progress_bar = txtProgressBar(min=0, max=2000, style = 1, char="=")
#
# for(i in 1:length(indel_dat$processid)){
# 	out_data = coi5p_pipe(indel_dat$test_seq[i],
# 							name = indel_dat$processid[i])
#
# 	indel_censored_translation_df = rbind(indel_censored_translation_df, flatten_coi5p(list(out_data)))
# 	setTxtProgressBar(progress_bar, value = i)
# }
# close(progress_bar)

write_csv(indel_censored_translation_df, "indel_data_censored_translation_output.csv")

###########################################
# messy, known translation tables
###########################################
messy_dat = read_csv(messy_seq_file)

messy_known_translation_data_output = mclapply(1:length(messy_dat$processid), function(i){
 coi5p_pipe(messy_dat$test_seq[i],
            name = messy_dat$processid[i],
            trans_table = messy_dat$genbankid[i])
},mc.cores = ncores)

messy_known_translation_df = flatten_coi5p(messy_known_translation_data_output)

#Alternative workflow for low memory machine:
# messy_known_translation_df = data.frame(matrix(ncol = 7, nrow = 0),stringsAsFactors = FALSE )
# colnames(messy_known_translation_df) = c("name", "raw", "framed", "aaSeq", "aaScore", "messy_likely", "stop_codons")
#
# progress_bar = txtProgressBar(min=0, max=2000, style = 1, char="=")
# for(i in 1:length(messy_dat$processid)){
# 	out_data = coi5p_pipe(messy_dat$test_seq[i],
# 							name = messy_dat$processid[i],
# 							trans_table = messy_dat$genbankid[i])
#
# 	messy_known_translation_df = rbind(messy_known_translation_df, flatten_coi5p(list(out_data)))
# 	setTxtProgressBar(progress_bar, value = i)
# }
# close(progress_bar)

write_csv(messy_known_translation_df, "messy_data_known_translation_output.csv")

###########################################
# messy, censored translation
###########################################

messy_censored_translation_data_output = mclapply(1:length(messy_dat$processid), function(i){
 coi5p_pipe(messy_dat$test_seq[i],
            name = messy_dat$processid[i])
},mc.cores = ncores)

messy_censored_translation_df = flatten_coi5p(messy_censored_translation_data_output)


#Alternative workflow for low memory machine:
# messy_censored_translation_df = data.frame(matrix(ncol = 7, nrow = 0),stringsAsFactors = FALSE )
# colnames(messy_censored_translation_df) = c("name", "raw", "framed", "aaSeq", "aaScore", "messy_likely", "stop_codons")
#
# progress_bar = txtProgressBar(min=0, max=2000, style = 1, char="=")
# for(i in 1:length(messy_dat$processid)){
# 	out_data = coi5p_pipe(messy_dat$test_seq[i],
# 							name = messy_dat$processid[i])
#
# 	messy_censored_translation_df = rbind(messy_censored_translation_df, flatten_coi5p(list(out_data)))
# 	setTxtProgressBar(progress_bar, value = i)
# }
# close(progress_bar)

write_csv(messy_censored_translation_df, "messy_data_censored_translation_output.csv")
```
