## supplementary for "coil: an R package for cytochrome C oxidase I (COI) DNA barcode data cleaning, translation, and error evaluation": S4-constrained_eval.r

```
# simulated metabarcode style sequence (non-full-length barcode) analysis with coil
library(coil)
library(tidyverse)

library(ROCit)
library(pROC)

#unzip Supplementary folder #6, and make that the working directory
setwd('S6-validation_data')

print("set the desired COI-5P sub-region here:")
start = 337
stop = 636

###############################################
#read in the supplementary data
###############################################

clean_seq_file = 'inputs/sequence_no_changes_2k.csv'
indel_seq_file = 'inputs/sequence_indel_2k.csv'

clean_dat = read_csv(clean_seq_file)
indel_dat = read_csv(indel_seq_file)

###############################################
#obtain relevant subsets of sequences from the data
###############################################

indel_dat$insert_vec = lapply(indel_dat$inserton_positions, 
	function(x){
		as.integer(strsplit(x, ";")[[1]])
		})

indel_dat$del_vec = lapply(indel_dat$deletion_positions, 
	function(x){
		as.integer(strsplit(x, ";")[[1]])
		})


err_in_range = function(pos_vec, start, stop){
	if(!is.na(pos_vec)){
		for(i in pos_vec){
			if (i > start & i < stop){
				return(TRUE)
			}
		}
	}
	return(FALSE)
}


indel_dat$insert_in_range = unlist(lapply(1:nrow(indel_dat), function(i){
	err_in_range(indel_dat$insert_vec[[i]], 350, 620)
	}))

indel_dat$delete_in_range = unlist(lapply(1:nrow(indel_dat), function(i){
	err_in_range(indel_dat$del_vec[[i]], 350, 620)
	}))

indel_dat$err_in_range = indel_dat$insert_in_range | indel_dat$delete_in_range

indel_dat$err_in_range

relevant_indel = indel_dat[indel_dat$err_in_range,]
nrow(relevant_indel)

#obtian a subset of the full dna sequences
subset_sequence = function(seq, start, stop){
	s_vec = strsplit(seq, "")[[1]]
	out_vec = s_vec[start:stop]
	return(paste(out_vec, collapse=""))
}

relevant_indel$sub_seq = unlist(lapply(1:nrow(relevant_indel), function(i){
	subset_sequence(relevant_indel$test_seq[[i]], start, stop)
	}))


clean_dat$sub_seq = unlist(lapply(1:nrow(clean_dat), function(i){
	subset_sequence(clean_dat$test_seq[[i]], start, stop)
	}))


###############################################
#define a constrained PHMMs to assess sequences with
###############################################

#337 - 636 is our simulated metabarcode (non-full-length barcode) region
#want to start at position 337 and cover 300bp
# you can change these values and rerun the subsequent script to get a suggested
# AUC cutoff for your own customized 
nt_start = start
nt_end = stop

#Get the corresponding amino acid start and end points
#the start and end positions are different than the nucleotide numbers, because 3bp make one amino acid
aa_start = ceiling(nt_start/3) # ceiling is used because 337/3 = 112.333, i.e. the first base pair of amino acid number 113
aa_end = ceiling(nt_end/3)
 
meta_nt_phmm = subsetPHMM(nt_coi_PHMM, start = nt_start, end = nt_end)
meta_aa_phmm = subsetPHMM(aa_coi_PHMM, start = aa_start, end = aa_end)


###############################################
#run the coil analysis on the subsequences, with the constrained PHMMs. known translation tables
###############################################

relevant_indel_outputs = lapply(1:nrow(relevant_indel), function(i){
	coi5p_pipe(relevant_indel$sub_seq[[i]], 
				name = relevant_indel$processid[[i]],
				trans_table = relevant_indel$genbankid[[i]],
				nt_PHMM = meta_nt_phmm, 
				aa_PHMM = meta_aa_phmm)	
	})


clean_dat_outputs = lapply(1:nrow(clean_dat), function(i){
	coi5p_pipe(clean_dat$sub_seq[[i]], 
				name = clean_dat$processid[[i]],
				trans_table = clean_dat$genbankid[[i]],
				nt_PHMM = meta_nt_phmm, 
				aa_PHMM = meta_aa_phmm)	
	})


###############################################
# label and bind the two dataframes, assess AUC
###############################################

relevant_indel_output_df = flatten_coi5p(relevant_indel_outputs)
relevant_indel_output_df$error_present = TRUE

clean_dat_output_df = flatten_coi5p(clean_dat_outputs)
clean_dat_output_df$error_present = FALSE

sum(relevant_indel_output_df$stop_codons)
sum(clean_dat_output_df$stop_codons)


met_barcode_sim = rbind(relevant_indel_output_df, 
								clean_dat_output_df)

aa_ROC = rocit(score=met_barcode_sim$aaScore, class=!met_barcode_sim$error_present)

class_measure =  measureit(score = met_barcode_sim$aaScore, class = !met_barcode_sim$error_present,
                     measure = c("ACC", "SENS", "SPEC", "FSCR"))

class_metrics = as.data.frame(cbind(Cutoff = class_measure$Cutoff, Depth = class_measure$Depth,
                                 Accuracy = class_measure$ACC, Sensitivity = class_measure$SENS,
                                 Specificity = class_measure$SPEC, FScore = class_measure$FSCR))

class_metrics = class_metrics[!is.na(class_metrics$FScore),]
optimal_cutoff = class_metrics$Cutoff[class_metrics$FScore == max(class_metrics$FScore, na.rm=TRUE)]


###############################################
# optimal cutoff and AUC
###############################################

print("optimal cutoff and AUC with known translation")


print("cutoff")
print(censored_optimal_cutoff) 
print("AUC")
print(aa_ROC$AUC)     # auc = 0.996 
summary(aa_ROC)
plot(aa_ROC)


met_barcode_sim_roc = roc(!met_barcode_sim$error_present, 
							met_barcode_sim$aaScore)

g_met_barcode_sim_roc = ggroc(met_barcode_sim_roc) + theme_minimal() + 
    geom_segment(aes(x = 1, xend = 0, y = 0, yend = 1), color="grey", linetype="dashed")+
    theme(axis.title=element_text(size=16, face='bold'),
    		axis.text=element_text(size=16), 
    		plot.title = element_text(size = 22))

ggsave('g_met_barcode_sim_roc.jpeg', g_met_barcode_sim_roc, width = 8, height = 8)


###############################################
#repeat the coil analysis on the subsequences, with the constrained PHMMs
#this time using the constrained models
###############################################
print("repeating the analysis using censored translation")

#no trans table passed so default val (censored translation) will be used
censored_relevant_indel_outputs = lapply(1:nrow(relevant_indel), function(i){
	coi5p_pipe(relevant_indel$sub_seq[[i]], 
				name = relevant_indel$processid[[i]],
				nt_PHMM = meta_nt_phmm, 
				aa_PHMM = meta_aa_phmm)	
	})


censored_clean_dat_outputs = lapply(1:nrow(clean_dat), function(i){
	coi5p_pipe(clean_dat$sub_seq[[i]], 
				name = clean_dat$processid[[i]],
				nt_PHMM = meta_nt_phmm, 
				aa_PHMM = meta_aa_phmm)	
	})


###############################################
# label and bind the two dataframes, assess AUC
###############################################

censored_relevant_indel_output_df = flatten_coi5p(censored_relevant_indel_outputs)
censored_relevant_indel_output_df$error_present = TRUE

censored_clean_dat_output_df = flatten_coi5p(censored_clean_dat_outputs)
censored_clean_dat_output_df$error_present = FALSE

sum(censored_relevant_indel_output_df$stop_codons)
sum(censored_clean_dat_output_df$stop_codons)


censored_met_barcode_sim = rbind(relevant_indel_output_df, 
								clean_dat_output_df)

aa_ROC = rocit(score=censored_met_barcode_sim$aaScore, class=!censored_met_barcode_sim$error_present)

class_measure =  measureit(score = censored_met_barcode_sim$aaScore, class = !censored_met_barcode_sim$error_present,
                     measure = c("ACC", "SENS", "SPEC", "FSCR"))

class_metrics = as.data.frame(cbind(Cutoff = class_measure$Cutoff, Depth = class_measure$Depth,
                                 Accuracy = class_measure$ACC, Sensitivity = class_measure$SENS,
                                 Specificity = class_measure$SPEC, FScore = class_measure$FSCR))

class_metrics = class_metrics[!is.na(class_metrics$FScore),]
censored_optimal_cutoff = class_metrics$Cutoff[class_metrics$FScore == max(class_metrics$FScore, na.rm=TRUE)]


###############################################
# optimal cutoff and AUC
###############################################
print("optimal cutoff and AUC with censored translation")

print("cutoff")
print(censored_optimal_cutoff) 
print("AUC")
print(aa_ROC$AUC)     
summary(aa_ROC)
plot(aa_ROC)


met_barcode_sim_roc = roc(!met_barcode_sim$error_present, 
							met_barcode_sim$aaScore)

g_met_barcode_sim_roc = ggroc(met_barcode_sim_roc) + theme_minimal() + 
    geom_segment(aes(x = 1, xend = 0, y = 0, yend = 1), color="grey", linetype="dashed")+
    theme(axis.title=element_text(size=16, face='bold'),
    		axis.text=element_text(size=16), 
    		plot.title = element_text(size = 22))

ggsave('g_met_barcode_sim_roc_censored_translation.jpeg', g_met_barcode_sim_roc, width = 8, height = 8)
```
