## supplementary for "coil: an R package for cytochrome C oxidase I (COI) DNA barcode data cleaning, translation, and error evaluation": S7-coil-vignette.pdf

*Cameron M. Nugent*

*2019-11-01*

```r
#install.packages('coil')
library(coil)
```

### Abstract

**coil** is an R package designed for the cleaning, contextualization, and assessment of cytochrome c oxidase I DNA barcode data (COI-5P, or the five prime portion of COI). It contains functions for placing COI-5P barcode sequences into a common reading frame, translating DNA sequences to amino acids, and for assessing the likelihood that a given barcode sequence includes an insertion or deletion error. These functions are provided as a single function analysis pipeline and are also available individually for efficient and targeted analysis of barcode data.

### Introduction

The backbone of the `coil` package is a pair of profile hidden Markov models (PHMMs) that have been trained using a representative sample of the COI-5P sequences available on the BOLD database. A 657 nucleotide PHMM receives raw sequences from the user and uses the Viterbi algorithm (implemented via the R package aphid) to match the input sequence against the COI-5P nucleotide profile. The second PHMM receives an amino acid sequence that is matched against the COI-5P amino acids profile. The model provides two Boolean output metrics to the user: (a) the sequence contains stop codons (T/F), (b) is the sequence likely to contain an insertion or deletion error (T/F). The insertion or deletion Boolean is based on the log likelihood of the amino acid sequence compared to the PHMM. A default indel likelihood threshold of -358.88 is set, but this can be changed by the user. Sequences with likelihood values less than this threshold indicate the sequence is likely to contain an indel error, as the amino acid sequence is improbable and therefore indicative of a possible frame shift.

The nucleotide and amino acid PHMMs are interfaced through the translate function, which takes the in-frame nucleotide sequence and translates it to amino acids. This function uses the sequinr package to conduct translation in all instances where the genetic code associated with the sample is known. For samples without taxonomic IDs or known genetic codes, an additional genetic code is provided. This genetic code is used to conduct censored translation, meaning that translation is conducted normally for codons that do not vary in the amino acid they code for across all known animal mitochondrial genetic codes. The codons that are known to vary in the amino acid they code for across taxa are not translated; rather a placeholder (`?`) is output to indicate that the amino acid at this location in the sequence cannot be stated with certainty. This functionality allows the `indel_check` function to assess the likelihood of sequences of unknown taxonomy without being overly stringent in its characterization of sequences as indels due to the appropriation of the wrong translation table.

```
Censored translation table:
           FFLLSSSSYY?*CCWWLLLLPPPPHHQQRRRRII?MTTTTNN?KSS??VVVVAAAADDEEGGGG
  Base1  = TTTTTTTTTTTTTTTTCCCCCCCCCCCCCCCCAAAAAAAAAAAAAAAAGGGGGGGGGGGGGGGG
  Base2  = TTTTCCCCAAAAGGGGTTTTCCCCAAAAGGGGTTTTCCCCAAAAGGGGTTTTCCCCAAAAGGGG
  Base3  = TCAGTCAGTCAGTCAGTCAGTCAGTCAGTCAGTCAGTCAGTCAGTCAGTCAGTCAGTCAGTCAG
```

The translation table employed in censored translation - five codons are translated to placeholder question marks, due to their ambiguity across different mitochondrial translation tables.

### The coil package

#### Dependencies

The `coil` package is dependent on the `aphid` package for comparison of sequences against the COI-5P PHMMs. The `ape` package is a requirement as well because `coil` internally converts all DNA and amino acid sequences to the ape "DNAbin" and "AAbin" object types to increase computational efficiency. As previously stated, `coil` is also dependent on the sequinr package for translation of sequences when the genetic code associated with the sample is known.

#### Using the package

##### Full analysis pipeline for a single sequence

An example execution of the complete coil analysis pipeline with default options is demonstrated below using an example COI-5P barcode DNA sequence.

```
output = coi5p_pipe(example_nt_string)
output
```

```
## coi5p barcode sequence
## raw sequence:
## ctctacttgatttttggtgcatgag...ggacccaattctctatcaacactta
## framed sequence:
## ---ctctacttgatttttggtgcat...ggacccaattctctatcaacactta
## Amino acid sequence:
## -LYLIFGAWAG?VG?ALSLLIRAEL...LTDRNLNTTFFDPAGGGDPILYQHL
## Raw sequence was trimmed: FALSE
## Stop codon present: FALSE, Amino acid PHMM score:-206.22045
## The sequence likely does not contain an insertion or deletion.
## Base pair 1 of the raw sequence is base pair 4 of the COI-5P region.
```

Executing the entire pipeline yields a coi5p object. Calling the variable name prints the coi5p object's summary and shows important information about the sequence.

Individual components can be obtained from the object using the dollar sign notation.

```
#see the available components
names(output)
```

```
##  [1] "name"         "raw"          "data"         "framed"
##  [5] "was_trimmed"  "align_report" "aaSeq"        "aaScore"
##  [9] "indel_likely" "stop_codons"
```

```
#retrieve only the amino acid sequence from the object
output$aaScore
```

```
## [1] -206.2204
```

By default the pipeline conducts censored translation, avoiding translation of the codons that are known to code for different amino acids among different species. If taxonomic information is available for the sample (available ranks: family, order, class, phylum), in most cases you can use the helper function `which_trans_table` to determine the proper genetic code to use. If the taxonomic group contains species that have different genetic codes, a 0 is returned to indicate that it is a good idea to use censored translation.

```r
ex_table_to_use = which_trans_table("Scyliorhinidae")
ex_table_to_use
```

```
## [1] 2
```

The analysis can then be run with a non-censored translation step. Note below that the amino acid sequence is now devoid of question marks and the PHMM score is lower.

```r
output = coi5p_pipe(example_nt_string, trans_table = ex_table_to_use)
output
```

```
## coi5p barcode sequence
## raw sequence:
## ctctacttgatttttggtgcatgag...ggacccaattctctatcaacactta
## framed sequence:
## ---ctctacttgatttttggtgcat...ggacccaattctctatcaacactta
## Amino acid sequence:
## -LYLIFGAWAGMVGMALSLLIRAEL...LTDRNLNTTFFDPAGGGDPILYQHL
## Raw sequence was trimmed: FALSE
## Stop codon present: FALSE, Amino acid PHMM score:-103.65363
## The sequence likely does not contain an insertion or deletion.
## Base pair 1 of the raw sequence is base pair 4 of the COI-5P region.
```

#### Calling functions individually

There are four functions that constitute the coi5p analysis pipeline: `coi5p`, `frame`, `translate` and `indel_check`. These are available to the user individually, for instances where only part of the analysis pipeline is needed (i.e. if you wish to frame sequences but not waste resources translating them, you could run only `coi5p` and `frame`).

```r
  #build the coi5p object
  dat = coi5p(example_nt_string, name = "example_sequence_1")
  #frame the sequence
  dat = frame(dat)
  #since we determined the genetic code above, we can use
  #the proper translation table as opposed to conducting
  #the default censored translation
  dat = translate(dat, trans_table = 2)
  #check to see if an insertion or deletion is likely
  dat = indel_check(dat)
  dat
```

```
## coi5p barcode sequence: example_sequence_1
## raw sequence:
## ctctacttgatttttggtgcatgag...ggacccaattctctatcaacactta
## framed sequence:
## ---ctctacttgatttttggtgcat...ggacccaattctctatcaacactta
## Amino acid sequence:
## -LYLIFGAWAGMVGMALSLLIRAEL...LTDRNLNTTFFDPAGGGDPILYQHL
## Raw sequence was trimmed: FALSE
## Stop codon present: FALSE, Amino acid PHMM score:-103.65363
## The sequence likely does not contain an insertion or deletion.
## Base pair 1 of the raw sequence is base pair 4 of the COI-5P region.
```

#### Example of a batch analysis of barcode sequences

Here we will be working with the example dataframe: `example_barcode_data`. Although loading and outputting DNA sequence data in R is outside of the scope of the `coil` package, the supplementary section at the end of this vignette includes an example of how one can load a fasta file into a dataframe with a structure matching that of `example_barcode_data`.

`example_barcode_data` contains 9 barcode sequences that demonstrate the different abilities of the `coil` package. Some sequences are longer than the barcode COI-5P barcode region, some are shorter, and some have insertion or deletion errors.

```
#this is the example data set
dim(example_barcode_data)
```

```
## [1] 9 5
```

```
names(example_barcode_data)
```

```
## [1] "id"           "genetic_code" "taxa"         "sequence"
## [5] "notes"
```

```
# to look at the full dataframe:
# example_barcode_data
```

The coi5p analysis pipeline can be applied to a dataframe in a iterative fashion. Here the pipeline is implemented through the use of the `lapply` function, which lets the unique sequence, id, and genetic code of each row in the dataframe be passed into the `coi5p_pipe` function.

```
example_barcode_data$coi_output = lapply(1:length(example_barcode_data$id), function(i){
  coi5p_pipe(example_barcode_data$sequence[i],
             name = example_barcode_data$id[i],
             trans_table = example_barcode_data$genetic_code[i])
})

example_barcode_data$coi_output[[1]] #example of the first output
```

```
## coi5p barcode sequence: ex_1_clean
## raw sequence:
## acgctttactttatttttggcatgt...taaccctattctttaccagcatttg
## framed sequence:
## acgctttactttatttttggcatgt...taaccctattctttaccagcatttg
## Amino acid sequence:
## TLYFIFGMWAGFIGLSMSLLIRMEL...LTDRNFNTSFFDPSGGGNPILYQHL
## Raw sequence was trimmed: FALSE
## Stop codon present: FALSE, Amino acid PHMM score:-184.28122
## The sequence likely does not contain an insertion or deletion.
## Base pair 1 of the raw sequence is base pair 1 of the COI-5P region.
```

Tip: to increase speed, use `mclapply` from the base R `parallel` package as opposed to `lapply`.

The coi5p objects are nested within the dataframe. Individual components can be extracted from the object as needed using the dollar sign notation. Below `lapply` is used to extract the framed sequence from each coi5p object and turn it into its own column in the dataframe. As we can see from the output below, dashes have been added to the front of the short sequence, if we compare the framed sequences from the long inputs (rows 5 and 6) to their original sequence, we see that coi5p has trimmed the sequence outside of the barcode region.

```
example_barcode_data$framed_seq = unlist(lapply(example_barcode_data$coi_output,
  function(x){
    x$framed
}))
```

```r
#has coi5p trimmed characters?
nchar(example_barcode_data$framed_seq[[5]]) < nchar(example_barcode_data$sequence[[5]])
```

```
## [1] TRUE
```

The `lapply` notation used above is rather clunky, so the coi5p package contains a helper function to aid the user in flattening a list of coi5p objects into a dataframe. By default, all of the available object components will be output to the dataframe, but the user can choose a subset of components they wish to extract from the object. Note: this function assumes that the coi5p objects in the list have been put through the same workflow and therefore have a matching set of components. As an example of how you could break it, if you've applied the `translate` function to only one member of the list and not the others then the coi5p objects will have non-matching sets of components and `flatten_coi5p` will not work properly.

```r
#extract only a single column
col_df = flatten_coi5p(example_barcode_data$coi_output, keep_cols = 'aaSeq')
#extract multiple columns
multi_df = flatten_coi5p(example_barcode_data$coi_output, keep_cols = c('framed','aaSeq'))
#extract all columns
full_coi5p_df = flatten_coi5p(example_barcode_data$coi_output)
#full_coi5p_df
```

The memory requirements of the method demonstrated above are trivial because the example dataframe has only nine rows. If millions of sequences are being processed, then keeping all of the coi5p objects in memory at once may become prohibitive (this is likely not an issue for most users). This will depend on the amount of RAM available on your machine. The average `coi5p_pipe` output is ~6KB in size, so processing 1 million sequences at once would occupy ~6GB of RAM. If you are trying to limit RAM usage, the following workflow can help keep RAM requirements modest by instantiating only one coi5p object at a time, but as a trade-off may take slightly more time to execute.

```r
full_coi5p_df = data.frame(matrix(ncol = 9, nrow = 0),stringsAsFactors = FALSE )
colnames(full_coi5p_df) = c("name", "raw", "framed", "was_trimmed", "align_report",
                            "aaSeq", "aaScore", "indel_likely", "stop_codons")

for(i in 1:length(example_barcode_data$id)){
    out_data = coi5p_pipe(example_barcode_data$sequence[i],
                          name = example_barcode_data$id[i],
                          trans_table = example_barcode_data$genetic_code[i])
  #for extreme memory conservation - could write each line of output to a .csv
    #instead of binding it to an output dataframe.
    full_coi5p_df = rbind(full_coi5p_df, flatten_coi5p(list(out_data)))
}
```

As demonstrated here, the `coil` package allows for robust cleaning, contextualization and error assessment of novel COI-5P barcode data. The package's analysis pipeline is designed in a modular fashion, allowing the user to run only the functions required for their given use case. The pipeline is designed with scalability in mind; each sequence is processed individually, allowing for parallelization to optimize analysis speed (i.e. via R's `mclapply` function) when computational resources are abundant or for the sequential analysis of sequences when limited memory is available.

#### Analysis of metabarcode data - Sub-setting the PHMM

The `coil` package's performance is sub-optimal when the length of the sequence being processed is shorter than the length of the PHMM it is being compared against (Nugent et al. 2019 doi: https://doi.org/10.1101/2019.12.12.865014). When sequences are shorter than the PHMM profile length, the inferred reading frame can be incorrect in up to 5% of sequences. This error occurs when the `frame` function matches the leading nucleotides of the given query to an incorrect position in the PHMM (a consequence of sequence similarity between different parts of the barcode region). This is especially problematic in the processing of metabarcoding data (i.e. using coil to error check consensus sequences of operational taxonomic units) because metabarcoding often targets shorter, standardized sections of the full barcode region.

To optimize performance for metabarcode data, `coil` contains a function, `subsetPHMM`, that allows for input sequences to be compared against a subset of the full COI-5P barcode region. This allows for `coil` to be effectively employed in the processing and error evaluation of metabarcode data, if the user knows which part of the COI-5P barcode region has been targeted.

To demonstrate this process, we will consider the following two ~300bp barcode fragments. These are derived from `coil`'s `example_nt_string` below. `dna_336_subset` is an error-free 300bp fragment, and `dna_336_subset_indel` is the same fragment with an deletion error introduced.

```
dna_vector = strsplit(example_nt_string, "")[[1]]
#three dashes added to the sequence because the example_nt_string starts at codon 2
dna_vector = c("-", "-", "-", dna_vector)
dna_336_subset = paste(dna_vector[336:635], collapse="")

#deleted a base pair from the sequence, simulating an indel error
dna_336_subset_indel = paste(c(dna_vector[336:358]  ,dna_vector[360:635]), collapse="")
```

Since `dna_336_subset` is much shorter than the full nucleotide PHMM (included in coil as the variable: `nt_coi_PHMM`), when the sequence is analyzed with the `coi5p_pipe` function, a false match occurs and the reading frame is incorrectly established. The sequence is reported to contain stop codons, when it is in fact a true 300bp barcode fragment. This is an example of why false positives are sometimes produced for shorter sequences.

```
false_pos = coi5p_pipe(dna_336_subset)
false_pos$stop_codons
```

```
## [1] TRUE
```

If we know the region of barcode that are barcode sequence fragment corresponds to, we can subset the nucleotide and amino acid PHMMs. Passing the subset models to the `coi5p_pipe` function along with the query sequence allows us to compare the sequence to only the region of interest.

To default PHMMs are `nt_coi_PHMM` and `aa_coi_PHMM`, which respectively are trained on nucleotide and amino acid sequences of the COI-5P region. To subset each PHMM, we pass the PHMM and the start and end positions of the sub-region of interest to the `subsetPHMM` function. This produces new models which we can compare smaller sequences against.

*IMPORTANT NOTE:* When sub-setting `nt_coi_PHMM` with `subsetPHMM` it is strongly advised that your `start` position is the first base pair of a codon (codon 1 starts at position 1, so the `start` argument should be part of the sequence 1,4,7,10,13,16,etc). The cell below contains a small function (`first_bp_of_codon`) to verify this. If you do not heed this advice, you will need to pass a `frame_offset` argument to the `coi5p_pipe` function (i.e. if your start value is `bad_start = 5`, the corresponding needs to be included in the `coi5p_pipe` function call `frame_offset = (bad_start-1)%%3`), and this makes things complicated.

```
#want to start at position 337 and cover 300bp
nt_start = 337
nt_end = 636

#Get the corresponding amino acid start and end points
#the start and end positions are different than the nucleotide numbers,
#because 3bp make one amino acid
# ceiling is used because 337/3 = 112.333, i.e. the first base pair of amino acid 113
aa_start = ceiling(nt_start/3)
```

```r
aa_end = ceiling(nt_end/3)

meta_nt_phmm = subsetPHMM(nt_coi_PHMM, start = nt_start, end = nt_end)
meta_aa_phmm = subsetPHMM(aa_coi_PHMM, start = aa_start, end = aa_end)

#Addendum to note IMPORTANT NOTE:
#This function can be used to check your start is the first bp of a codon:
first_bp_of_codon = function(x){
 if(((x-1)%%3) == 0){
   return(TRUE)
 }
 return(FALSE)
}

first_bp_of_codon(nt_start)
```

```
## [1] TRUE
```

Now that we have our nucleotide and amino acid PHMMs representing the 300bp subset of the full barcode region, we can run the `coi5p_pipe` function, this time passing the query sequence along with our non-default PHMMs.

```r
#pass the dna sequence fragment with no error, and also subset the nt and aa PHMMs
subset_no_error_output = coi5p_pipe(dna_336_subset,
                                    nt_PHMM = meta_nt_phmm,
                                    aa_PHMM = meta_aa_phmm)
#is there evidence of stop codons:
subset_no_error_output$stop_codons
```

```
## [1] FALSE
```

```r
#see the full output
subset_no_error_output
```

```
## coi5p barcode sequence
## raw sequence:
## atatcctcctttagcaggtaattta...ttgaccccgcagggggaggggaccc
## framed sequence:
## tatcctcctttagcaggtaatttag...ctttgaccccgcagggggaggggac
## Amino acid sequence:
## YPPLAGNLAHAGPSVDLAIFSLHLA...AAGIT?LLTDRNLNTTFFDPAGGGD
## Raw sequence was trimmed: TRUE
## Stop codon present: FALSE, Amino acid PHMM score:-87.41969
## The sequence likely does not contain an insertion or deletion.
## Base pair 1 of the raw sequence is base pair 1 of the COI-5P region.
```

As we can see the error-free barcode sequence fragment has this time been framed properly, and as a result there is no evidence of a stop codon and the amino acid PHMM score is high. Below the same subset PHMMs are used to assess the barcode fragment with an indel error, it effectively identifies the presence of stop codons in the sequence (There the amino acid PHMM score is also low). Therefore using the `subsetPHMM` function allows us to effectively extend the functionality of `coil` to barcode sequence fragments by constraining the PHMMS. The frequency of false positives can be reduced and we can effectively separate barcode sequence fragments with indels from error-free sequences.

```r
subset_has_error_outpt = coi5p_pipe(dna_336_subset_indel,
                                    nt_PHMM = meta_nt_phmm,
                                    aa_PHMM = meta_aa_phmm)
```

```r
subset_has_error_outpt$stop_codons
```

```
## [1] TRUE
```

```r
subset_has_error_outpt
```

```
## coi5p barcode sequence
## raw sequence:
## atatcctcctttagcaggtaattag...ttgaccccgcagggggaggggaccc
## framed sequence:
## tatcctcctttagcaggtaattagc...ctttgaccccgcagggggaggggac
## Amino acid sequence:
## YPPLAGN*HMLAPLLI*PSFPFIWP...QPELQYY?QTATSTLHSLTPQGEG-
## Raw sequence was trimmed: TRUE
## Stop codon present: TRUE, Amino acid PHMM score:-491.58798
## The sequence likely contains an insertion or deletion.
## Base pair 1 of the raw sequence is base pair 1 of the COI-5P region.
```

#### Supplementary Information

##### Loading and manipulating a fasta file

The example of processing batch DNA barcode data above begins with the data in a clean dataframe. Since barcode data is not always obtained in a tidy format, some preprocessing by the user will likely be required. The following is provided to aid the user in developing a workflow for loading their barcode sequence data into R and constructing a tidy dataframe before beginning sequence analysis.

The example presented below shows how one can load a fasta file containing DNA sequences into R and then convert the sequence and header data into a tidy dataframe object. More information on the `read.fasta` function can be found in the seqinr documentation.

Information found in the header lines of fasta files varies, so the reader will likely need to alter this code for novel data sources. In this example, the header contains four fields (id, genetic code, taxa and notes) separated by a pipe character (`|`). The code below turns this fasta file into a dataframe that matches the `example_barcode_data` file used above.

```r
library(seqinr)
```

```
##
## Attaching package: 'seqinr'

## The following object is masked from 'package:coil':
##
##     translate
```

```r
# load the example fasta file included with coil
# included in the file's header line:
# the name of the sample, its genetic code, taxonomic designation and some notes
ex_fasta_file = system.file("extdata/example_barcode_data.fasta", package = "coil")

#read in the example fasta file using seqinr
ex_data = seqinr::read.fasta(ex_fasta_file, as.string = TRUE)

#here is what the output from read.fasta looks like
#head(ex_data)

#parse the data in the header line by splitting the name on the | character
```

```r
parsed_names_data = lapply(1:length(ex_data), function(i){
  unlist(strsplit(names(ex_data)[[i]],"\\|"))
})

# subset the components of the header line and build these and the sequence
# into a dataframe matching the style used in the coi5p batch example
example_barcode_data_from_scratch = data.frame(
  id = sapply(parsed_names_data, function(x) x[[1]]),
  genetic_code = sapply(parsed_names_data, function(x) x[[2]]),
  taxa = sapply(parsed_names_data, function(x) x[[3]]),
  sequence = unname(unlist(ex_data)),
  notes = sapply(parsed_names_data, function(x) x[[4]])
)

#uncomment the following line to see result
#head(example_barcode_data_from_scratch)
```
